# The ER membrane chaperone Shr3 acts in a progressive manner to assist the folding of related plasma membrane transport proteins

**DOI:** 10.1101/2020.07.05.188557

**Authors:** Ioanna Myronidi, Andreas Ring, Per O. Ljungdahl

**Author notes:** Corresponding author. Mailing address: SciLifeLab, Box 1031, SE-171 21 Solna, Sweden. Contributed equally, the names are listed in alphabetic order (Ioanna Myronidi, Andreas Ring).

## Abstract

Proteins with multiple membrane-spanning segments (MS) co-translationally insert into the endoplasmic reticulum (ER) membrane of eukaryotic cells. In *Saccharomyces cerevisiae*, Shr3 is an ER membrane-localized chaperone that is specifically required for the functional expression of amino acid permeases (AAP), a family of eighteen transporters comprised of 12 MS. Here, comprehensive scanning mutagenesis and deletion analysis of Shr3, combined with a modified split-ubiquitin approach, were used to probe chaperone-substrate interactions with seven different AAP *in vivo*. Our findings indicate that Shr3 specifically recognizes AAP substrates, largely independent of sequence-specific interactions involving membrane and luminally oriented domains. Shr3 selectively and robustly interacts with nested C-terminal AAP truncations in marked contrast to similar truncations of non-Shr3 substrate polytopic sugar transporters. Strikingly, Shr3-AAP interactions initiate with the first 4 MS of AAP and successively strengthen, but abruptly weaken when all 12 MS are present. The data are consistent with Shr3 acting in a temporal manner as a scaffold preventing AAP translation intermediates from engaging in non-productive interactions.

## Introduction

The integration and concomitant folding of membrane proteins in the lipid bilayer of the endoplasmic reticulum (ER) are critical steps in the biogenesis of transport proteins destined to function at the plasma membrane (PM). Most eukaryotic membrane proteins are cotranslationally inserted into the ER membrane via the Sec61-complex, also known as the translocon. The translocon forms a protein-conducting channel that mediates protein translocation and the co-translational partitioning of membrane-spanning segments (MS) (Johnson and van Waes, 1999; Rapoport et al., 2017; Seinen and Driessen, 2019). During the synthesis of complex polytopic membrane proteins comprised of multiple MS, each MS sequentially exits the channel and partitions into the ER membrane via the lateral gate of the translocon. Although the central channel of the translocon is too small to accommodate multiple MS, the translocon appears to have a limited capacity to promote the folding of membrane proteins comprised of up to three MS; two separate extra-channel MS-binding sites have been reported to act in a chaperone-like manner to delay the release of N-terminal MS until translation is completed (Hou et al., 2012). However, as the complexity of membrane proteins grows beyond three MS, the challenge of preventing inappropriate interactions between incompletely translated nascent chains apparently exceeds the chaperone-like activity of the translocon. Specifically, during the translation of complex polytopic membrane proteins, e.g., amino acid permeases with 12 MS, the N-terminal MS partition into the membrane prior to the synthesis and partitioning of C-terminal MS. To prevent the MS of translation intermediates from entering non-productive folding pathways, discrete and highly specialized ER resident membrane proteins have been described in fungi that prevent misfolding of specific families of polytopic membrane proteins (Kota and Ljungdahl, 2005; Lau et al., 2000; Ljungdahl et al., 1992; Luo et al., 2002; Martínez and Ljungdahl, 2000; 2004; Sherwood and Carlson, 1999; Shurtleff et al., 2018).

Shr3, the most comprehensively studied of these specialized ER components, was identified as an integral membrane protein required for the functional expression of the conserved family of amino acid permeases (AAP) (Ljungdahl *et al*., 1992). The AAP family in *Saccharomyces cerevisiae*, belonging to the Amino acid-Polyamine-Organocation (APC) super-family of transporters (Gilstring and Ljungdahl, 2000; Jack et al., 2000; Saier, 2000; Wong et al., 2012), is comprised of 18 genetically distinct but structurally similar proteins with 12 MS. Shr3 is composed of 210 amino acids organized into two functional domains; a N-terminal membrane domain comprised of four hydrophobic α-helices connected by three hydrophilic loops and a hydrophilic cytoplasmically oriented C-terminal domain. Initially, Shr3 was recognized as an essential factor facilitating the packaging of AAP into ER-derived secretory vesicles (Kuehn et al., 1996), and it was found to be important for the proper presentation of ER-exit motifs, located within the hydrophilic C-terminal tails of AAP, to the inner COPII coatomer subunit Sec24 (Kuehn et al., 1998; Malkus et al., 2002; Miller et al., 2002; Miller et al., 2003). Additional data regarding the packaging activity of Shr3 included genetic and physical interactions of Shr3 with components of the COPII-coated vesicles (Gilstring et al., 1999). Also, it was shown that Shr3 facilitated ER-vesicle formation in close proximity to fully integrated and folded AAP (Gilstring and Ljungdahl, 2000; Gilstring *et al*., 1999) and hence Shr3 was designated a packaging chaperone. The ability of Shr3 to interact with COPII components is primarily linked to its hydrophilic C-terminal tail. Consistently, Shr3 was found to associate with newly synthesized Gap1 in a transient manner, the interaction was reported to exhibit a half-life of approximately 15 min. The observed association with Gap1 was initially thought to occur post-translationally since Gap1 fully integrates into the ER-membrane with each MS correctly oriented independently of Shr3 (Gilstring and Ljungdahl, 2000). Also, the AAP that accumulate in the ER of *shr3*Δ strains do not activate the unfolded protein stress response (UPR) (Gilstring *et al*., 1999).

However, subsequent studies revealed that Shr3 has an important function that is separate from and precedes its packaging function. Consequently, the view of Shr3 evolved from being a packaging into a specialized membrane-localized chaperone that interacts early with substrate AAP during their co-translational insertion into the ER membrane (Kota et al., 2007; Kota and Ljungdahl, 2005). In these studies, Shr3 was found to prevent the aggregation of AAP in the ER-membrane, a function associated with its N-terminal membrane domain (Kota and Ljungdahl, 2005). Critical evidence demonstrating the importance of Shr3 in facilitating the folding of AAP includes the finding that co-expressed split N-and C-terminal portions of Gap1 assemble into a functional permease in a Shr3-dependent manner (Kota *et al*., 2007). The membrane domain of Shr3 is required and suffices to prevent aggregation of the first five MS of Gap1 enabling productive folding interactions with the C-terminal portions of Gap1. Similar to full-length Gap1, the N-terminal fragment displays an increased propensity to aggregate in membranes isolated from cells lacking Shr3, and importantly, its aggregation appeared not to be affected by the presence or absence of the C-terminal fragment. In marked contrast, the aggregation status of the C-terminal fragment was dependent on the presence of both Shr3 and the N-terminal fragment. Since the N- and C-terminal fragments individually insert into the membrane, Shr3 apparently can maintain the N-terminal fragment in a conformation that enables the C-terminal fragment to interact and assemble with it. Although direct physical interactions between Shr3 and substrate AAP have not been demonstrated, all available data is consistent with Shr3 interacting early with N-terminal MS as they co-translationally partition into the ER membrane.

During translocation, exclusively hydrophobic MS partition readily into the lipid phase of the membrane, whereas less hydrophobic MS containing (Heinrich et al., 2000) charged or polar residues partition into the membrane less readily and are retained in proximity to the translocon or to translocon associated proteins, e.g., TRAM (Heinrich and Rapoport, 2003). In analogy to TRAM, we posited that Shr3 facilitates the partitioning of MS of AAP containing charged or polar amino acid residues as they emerge from the translocon. According to this hypothesis, Shr3 may physically shield charged or polar residues within MS, thereby preventing these thermodynamically challenging segments from engaging in nonproductive interactions (Kota *et al*., 2007). More recent findings in yeast regarding the conserved ER membrane protein complex (EMC) have been interpreted in a similar manner (Miller-Vedam et al., 2020; Shurtleff *et al*., 2018). However, a striking difference between Shr3 and EMC function is that null alleles of *SHR3* do not activate the UPR (Gilstring *et al*., 1999).

Despite the clear requirement of Shr3 in AAP biogenesis, we currently lack critical information regarding the mechanisms underlying Shr3 function. Here we have focused on the membrane domain of Shr3 and employed a comprehensive scanning mutagenesis approach to define amino acid residues involved in recognizing AAP substrates. Further, we have exploited a split-ubiquitin approach to directly probe and characterize interactions with seven different AAP substrates *in vivo*. The data support Shr3 acting as a MLC providing a scaffold-like structure to help nascent chains of partially translated AAP maintain a structure required to enter and follow a productive folding pathway as translation proceeds to completion.

## Results

### Systematic scanning mutagenesis of MS within the membrane domain of Shr3

We have previously shown that the membrane domain of Shr3 is required and sufficient for facilitating the folding of AAPs (Kota and Ljungdahl, 2005). Here, a systematic scanning mutagenesis approach was used to identify residues within the Shr3 membrane domain required for function. Intramembrane residues were mutated to leucine; the length of consecutive substitution mutations varied, ranging from 2 to 13 residues. The extramembrane residues within ER lumenal loops L1 and L3 and cytoplasmic oriented NT and loop L2 where mutated to alanine; the length of consecutive alanine replacements ranged from 2 to 3.

The biological activity of the mutant proteins was initially assessed using growth-based assays on YPD supplemented with metsulfuron-methyl (MM), which provides a sensitive measure of Shr3 function. MM targets and inhibits branched-chain amino acid synthesis and growth is strictly dependent on the combined activity of multiple SPS-sensor regulated AAP that facilitate high-affinity isoleucine, leucine and valine uptake (Andréasson and Ljungdahl, 2002; Jørgensen et al., 1998). Serial dilutions of cell suspensions from strain JKY2 (*shr3*Δ) carrying vector control (VC), *SHR3* or the *shr3* mutant alleles were spotted on YPD and YPD+MM plates (Supplementary Material Fig. S1 – S10). Only three mutant alleles, *shr3-35, shr3-50* and *shr3-76*, failed to support growth (Fig. 1A and B). The steady state levels of the three mutant proteins were similar to wildtype Shr3 (Fig. 1B), suggesting that the mutant proteins were not grossly misfolded, and consequently, not prematurely targeted for ER-associated degradation.

**Figure 1.**
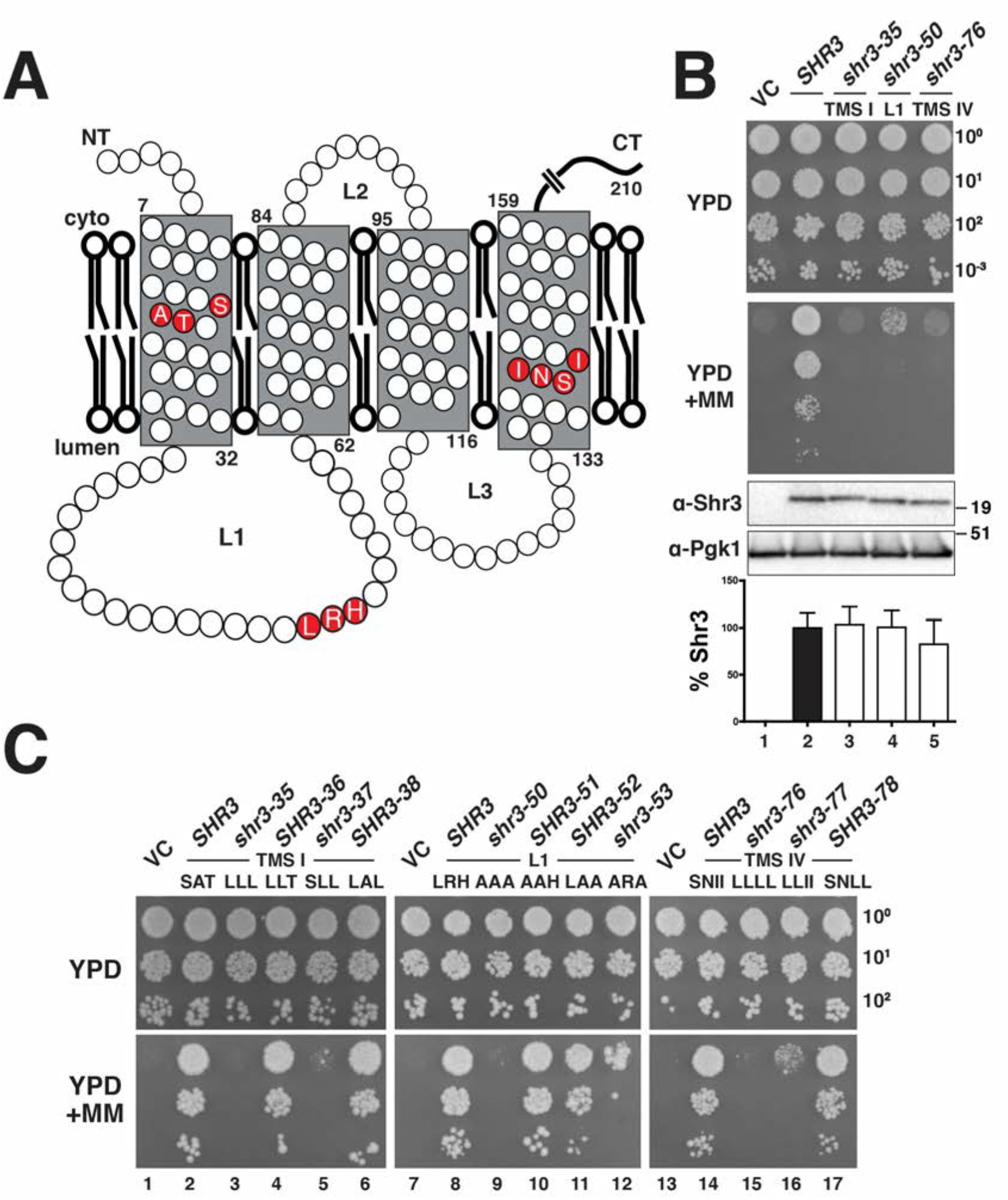
Scanning mutagenesis of the Shr3 membrane domain. (A) Graphical representation of Shr3 topology and position of residues resulting in a non-functional protein. (B) Top: Serial dilutions of cell suspensions from strain JKY2 (*shr3*Δ) carrying pRS316 (VC), pPL210 (*SHR3*), pAR4 (*shr3-35*), pAR18 (*shr3-50*) or pPL1349 (*shr3-76*) spotted on YPD and YPD+MM. The plates were incubated at 30 °C for 2 d and photographed. Bottom: Immunoblot analysis of Shr3 proteins in extracts prepared from the strains; the levels of Pgk1 were used as loading controls. The blots were developed using α-Shr3 and α-Pgk1 antibodies. The signal intensities of the immunoreactive forms of Shr3 and Pgk1 were quantified, and the Shr3 signals were normalized with respect to Pgk1; the mean values are plotted, error bars show standard deviation (n=3). (C) Serial dilutions of cell suspensions from strain JKY2 (*shr3*Δ) carrying pRS316 (VC), pPL210 (*SHR3*), pAR4 (*shr3-35*), pPL1330 (*SHR3-36*), pAR47 (*shr3-37*), pAR37 (*SHR3-38*), pAR18 (*shr3-50*), pAR51 (*SHR3-51*), pAR52 (*SHR3-52*), pAR50 (*shr3-53*), pPL1349 (*shr3-76*), pAR48 (*shr3-77*) or pAR49 (*SHR3-78*) spotted on YPD and YPD+MM plates. Plates were incubated at 30 °C for 2 d and photographed.

The *shr3-35* allele encodes a protein with residues 17 through 19 (serine-alanine-threonine) in MS I replaced by leucine (Fig. 1A). To more precisely define the critical residues, we constructed additional mutant alleles with paired leucine substitutions at residues 17-18 (LLT), 18-19 (SLL), and 17 and 19 (LAL). Expression of *SHR3-36* (LLT) and *SHR3-38* (LAL) alleles complemented *shr3*Δ and supported wildtype growth on YPD + MM (Fig. 1C, dilutions 4 and 6). By contrast, cells expressing *shr3-37* (SLL) did not complement, exhibiting a phenotype similar to the *shr3-35* (LLL) mutant (Fig. 1C, dilutions 3 and 5).

The *shr3-76* allele carries leucine replacements at residues 139 through 142 (serine-asparagine-isoleucine-isoleucine) in MS IV (Fig. 1A). Again, the importance of the affected residues was tested by creating alleles with paired leucine substitutions at positions 139-140 (LLII) and 141-142 (SNLL). Cells expressing *shr3-77* (LLII) allele grew poorly, although slightly better than cells expressing the *shr3-7*6 allele (Fig. 1C, dilutions 15 and 16). Cells expressing *SHR3-78* (SNLL) grew as wildtype *SHR3* (Fig. 1C, dilutions 14 and 17).

The third non-functional allele, *shr3-50*, encodes a mutant protein with alanine substitutions at residues 51 through 53 (leucine-arginine-histidine) located within the ER lumenal loop L1 (Fig. 1A). The importance of these residues was tested by paired alanine substitutions of the positions 51-52 (AAH), 52-53 (LAA), and 51 and 53 (ARA). Expression of these alleles demonstrated that *SHR3-51* (AAH) and *SHR3-52* (LAA) complemented *shr3*Δ similar to wildtype *SHR3* (Fig.1C, dilutions 8, 10 and 11). The *shr3-53* (ARA) allele exhibited reduced growth (Fig. 1C, dilution 12), indicative of compromised function. The finding that mutations affecting residues 53-55 in the ER lumenal loop L1 abolish function suggests that extramembrane sequences are important for guiding the folding of AAP sequences destined to be oriented towards the extracellular side of the PM.

### Deletion analysis of ER-lumen oriented loops

The possibility that Shr3 engages and interacts with its substrate AAP through contacts with extramembrane sequences prompted us to specifically test the functional significance of loops L1 and L3 (Fig. 2A). The residues 44-57 within L1 are predicted to fold into an amphipathic α-helix (Fig. 2B). We constructed four internal deletions in L1 that affect this secondary structure motif to varying extent: *shr3*Δ*90* (Δ34-48); *shr3*Δ*91* (Δ39-47); *shr3*Δ*92* (Δ44-54); and *shr3*Δ*93* (Δ55-60). Also, an internal deletion in L3 was constructed: *SHR3*Δ*94* (Δ121-127). The five deletion alleles directed the expression of mutant proteins at levels comparable to *SHR3* (Fig. 2C); the deletions do not decrease the steady state levels of protein. The function of these deletion alleles was assessed as before using YPD + MM. We also extended the analysis with more nuanced growth-based assays capable of monitoring amino acid uptake catalyzed predominantly by a single or a couple of AAP. This was accomplished by examining growth on minimal media individually supplemented with toxic amino acid analogues D-histidine, L-canavanine, and azetidine-2-carboxylate (AzC), which are taken up by Gap1 (Gresham et al., 2010), Can1 (Ono et al., 1983), Agp1/Gnp1 (Andréasson et al., 2004), respectively. The expression of functional alleles of *SHR3* results in impaired growth in the presence of these toxic analogues. Serial dilutions of cell suspensions from strain JKY2 (*shr3*Δ) carrying vector control (VC), *SHR3, shr3*Δ*90, shr3*Δ*91, shr3*Δ*92, shr3*Δ*93* or *SHR3*Δ*94* were spotted on SAD containing D-histidine, SD + L-canavanine, SD + AzC, and YPD + MM plates. The four internal deletion alleles affecting L1 failed to complement *shr3*Δ; the strains grew similar as the VC (Fig. 2C, dilutions 1, 3, 4, 5 and 6). By contrast, the strain expressing the internal deletion in L3 showed a more complex pattern of growth. On SAD + D-histidine and SD + L-canavanine, *SHR3*Δ*94* appeared to express a non-functional protein, the strain grew in the presence of these toxic amino acid analogues (Fig. 2C, compare dilution 1 with 7). However, on SD + AzC and YPD + MM, the *SHR3*Δ*94* allele exhibited growth similar to wildtype *SHR3* (Fig. 2C, compare dilution 2 with 7). Note that on YPD + MM the strain carrying *SHR3*Δ*94* exhibited enhanced growth compared to wildtype, and consequently, on YPD + MM we designated the phenotype WT^+^.

**Figure 2.**
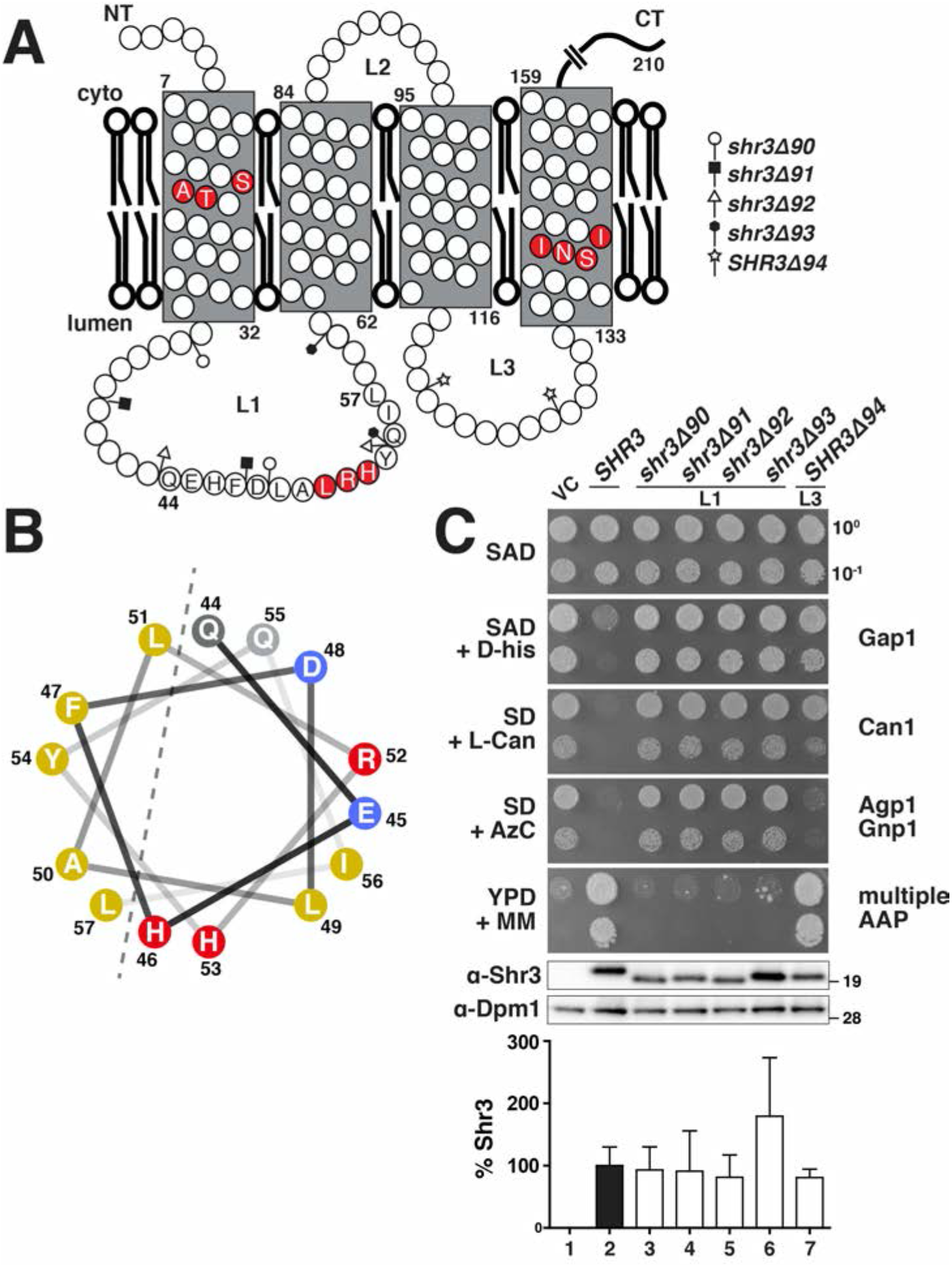
Deletion analysis of ER-lumen oriented loops. (A) Graphical representation of Shr3 topology and the positions of the internal deletions in loops L1 and L3. (B) Based on structural predictions (Drozdetskiy et al., 2015), amino acid residues 44-57 in L1 are predicted to fold into an α-helix with amphipathic character. Helical wheel projection of the L1 α-helix with non-polar (yellow), polar (grey), negatively-(blue) and positively-charged (red) residues indicated. (C) Serial dilutions of cell suspensions from strain JKY2 (*shr3*Δ) carrying pRS316 (VC), pPL210 (*SHR3*), pAR41 (*shr3*Δ*90*), pAR42 (*shr3*Δ*91*), pAR43 (*shr3*Δ*92*), pAR44 (*shr3*Δ*93*) or pAR45 (*shr3*Δ*94*) spotted on SAD containing D-histidine (D-his), SD + L-canavanine (L-Can), SD + AzC, and YPD + MM plates. Plates were incubated at 30 °C for 2 d and photographed. Bottom: Immunoblot analysis of Shr3 proteins in extracts prepared from the strains; the levels of Dpm1 were used as loading controls. The blots were developed using α-Shr3 and α-Dpm1 antibodies. The signal intensities of the immunoreactive forms of Shr3 and Dpm1 were quantified, and the Shr3 signals were normalized with respect to Dpm1; the mean values are plotted, error bars show standard deviation (n=3).

### Lumenal loop L3 influences substrate specificity

The finding that *SHR3*Δ*94* exhibited a range of phenotypes, i.e., from null to apparently enhanced functionality, prompted us to reexamine the growth characteristics of the 44 leucine- and alanine-scanning mutant alleles using the more nuanced growth-based assays. Serial dilutions of cell suspensions from strain JKY2 (*shr3*Δ) carrying vector control (VC), *SHR3* or one of the individual mutant alleles were spotted on SAD + D-histidine, SD + L-canavanine, SD + AzC and YPD+MM (Supplementary Material Fig. S1-S10). The growth characteristics were evaluated and the results, including the internal loop deletions, are summarized in an ordered heat-map (Fig. 3A). As was found in the initial evaluation of growth on YPD+MM, most of the mutant alleles were judged to encode functional proteins; the strains grew similarly as the strain carrying the *SHR3* wildtype control. Reevaluation of the three non-functional alleles, *shr3-35, shr3-50* and *shr3-76*, confirmed that the mutations exhibit major defects on all of the selective media (Fig. 3A). However, in some instances, several of the mutations, similar to *SHR3*Δ*94*, conferred robust growth on YPD+MM but did not complement *shr3*Δ on the other selective media. Strikingly, *SHR3-45, -63, -65, -68, -71, -74*, and *-75*, supported more robust growth on YPD + MM than *SHR3*, but exhibited a null phenotype on media containing toxic amino acid analogues. In summary, growth in the presence of D-histidine was found to be the most sensitive monitor of mutations in *SHR3*, perhaps due to the fact that D-amino acids are taken up by a single AAP, Gap1 (Grenson et al., 1970; Rytka, 1975). We note that mutations localized to the MS III and IV and the ER lumenal oriented loop L3 exhibited the most pleiotropic affects, suggesting that these regions of Shr3 facilitate interactions with discrete AAP, and potentially, comprise substrate specific determinants.

**Figure 3.**
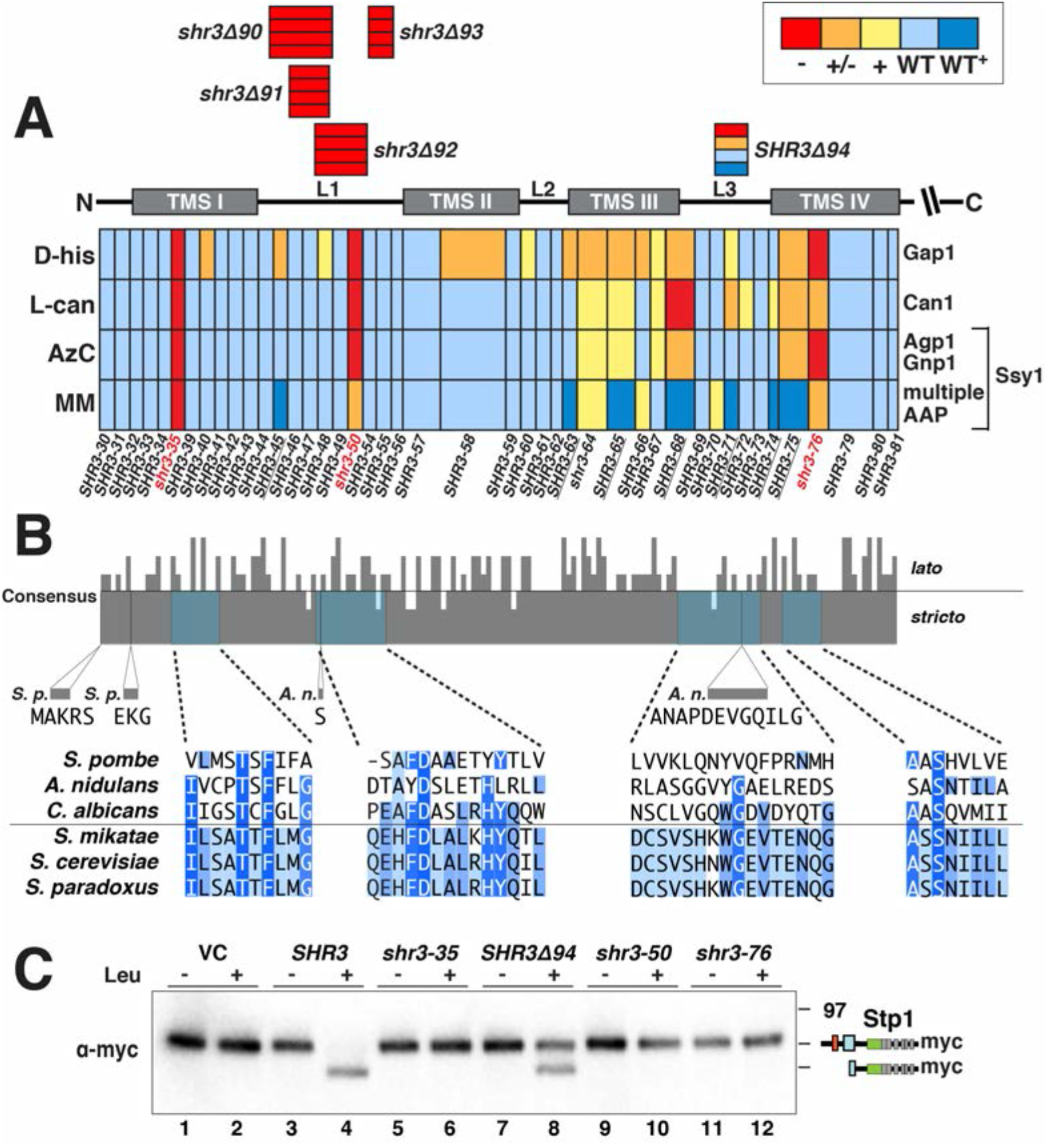
Mutational analysis of Shr3 function and substrate specificity. (A) Summary of growth characteristics of JKY2 (*shr3*Δ) individually expressing 44 Shr3-mutant proteins. Cells were spotted on media containing toxic amino analogues and nitrogen sources as follows: D-his, D-histidine (0.5% w/v), allantoin; L-can, L-canavanine (1 µg/ml), ammonium; AzC, azetidine-2-carboxylate (1 mM), ammonium; MM (200 µg/ml), yeast extract and peptone. Growth was scored after 2 - 3 d of incubation at 30 °C (Supplementary Materials Fig. S1-S10). Colors reflect Shr3 function relative to wildtype activity: red, no function (-); orange, weak but detectable function (+/-); yellow, intermediate function but less than wildtype (+); light blue, wildtype function (WT); dark blue, enhanced function (WT+). (B) Clustal O (Madeira *et al*., 2019) comparison of Shr3 sequences, corresponding to aa residues 1-159 of *S. cerevisiae*, and orthologs of members from the *Saccharomyces sensu stricto* group (*S. paradoxus, S. mikatae*) and orthologs from *sensu lato* fungi (*S. pombe, A. nidulans*, and *C. albicans*). The consensus plot (identity; (Waterhouse et al., 2009) and detailed multiple sequence alignments are presented for the regions with mutations giving rise to major growth defects on selective media; identical residues in three (light blue), four (blue), and five or six homologs (dark blue) are highlighted. (C) Shr3-dependent Ssy1 folding and function assessed by Stp1 processing. Immunoblot analysis of extracts from FGY135 (*shr3*Δ) carrying pCA204 (*STP1-13xMYC*) and pRS316 (VC), pPL210 (*SHR3*), pAR004 (*shr3-35*), pAR45 (*SHR3*Δ*94*), pAR018 (*shr3-50*) or pPL1351 (*shr3-76*). Cells were grown in SD and induced 30 min with 1.3 mM leucine (+) as indicated.

We performed multiple sequence alignments of the membrane domain of Shr3 and orthologs from two *Saccharomyces sensu stricto* strains, *S. paradoxus* and *S. mikatae*, and from three divergent *lato* fungal strains, *Candida albicans* (Csh3), *Schizosaccharomyces pombe* (Psh3) and *Aspergillus nidulans* (ShrA) and obtained a consensus identity plot (Fig. 3B) (Madeira et al., 2019). The Shr3 orthologs of *C. albicans, S. pombe* and *A. nidulans* have been shown to function analogously and are required for proper amino acid uptake. Heterologous expression of *CSH3* complements *shr3*Δ (Martínez and Ljungdahl, 2004), whereas heterologous expression of *PSH3* or *SHRA* only partially complement *shr3*Δ, merely facilitating the functional expression of limited subset of permeases (Erpapazoglou et al., 2006; Martínez and Ljungdahl, 2000). The Shr3 sequence is well-conserved in the *Saccharomyces sensu stricto* strains, exhibiting almost absolute identity. Several positions throughout the membrane domain of Shr3 are conserved between the full set of selected sequences. Interestingly, threonine 19, which is the single critical amino acid residue in MS I, is conserved in all orthologs (Fig. 3B, T19 is highlighted in dark blue). The requirement for a polar amino acid in MS IV of Shr3 is conserved as well (Fig. 3B, S139 highlighted in dark blue). A higher sequence divergence is evident in the luminal loop L3, with the extreme case of the *A. nidulans* orthologue that contains an extra sequence of twelve amino acid residues. The limited sequence identity in loop L3 aligns with the observation that mutations in L3 of Shr3 exhibit the most pleiotropic effects, and is consistent with the notion that L3 affects substrate specificity.

The finding that the Shr3Δ94 mutation supported robust growth on YPD+MM, suggested that the mutant protein retained the capacity to facilitate Ssy1 folding. In contrast to the other members of the AAP transporter family, Ssy1 functions as the primary receptor of extracellular amino acids in the context of the plasma membrane-localized SPS sensor (Didion et al., 1998; Iraqui et al., 1999; Klasson et al., 1999). In response to extracellular amino acids, Ssy1 initiates signaling events leading to the proteolytic activation of Stp1, which in turn induces the expression of several AAP genes including *AGP1* and *GNP1* and multiple permeases facilitating branched amino acid uptake. As an indirect measure of Shr3-Ssy1 interactions, we examined the proteolytic cleavage of the transcription factor Stp1 (Fig. 3C). Consistent with the growth assays, leucine induction led to Stp1 processing in strain FGY135 (*shr3*Δ) expressing *SHR3* or *SHR3*Δ*94* (Fig. 3C, lanes 4 and 8), but not the non-functional alleles *shr3-35, shr3-50* or *shr3-76* (Fig. 3C, lanes 6, 10 and 12).

### Shr3-AAP substrate interactions

To test if the observed growth phenotypes of the mutated *SHR3*/*shr3* alleles correlated with the ability of mutant Shr3/shr3 proteins to interact with specific AAP and facilitate their functional expression we exploited a split-ubiquitin approach to monitor Shr3-AAP interactions *in vivo* (Fig. 4A and 4B). A sequence encoding the N-terminal fragment of ubiquitin carrying the I13A mutation (NubA), which reduces the propensity of non-specific interactions (Johnsson, 2002; Johnsson and Varshavsky, 1994), was fused at the C-terminal end of *SHR3, shr3-35*, and *SHR3*Δ*94*, creating the Shr3-NubA constructs schematically depicted in Fig. 4A. Next, we created a *GAP1* allele encoding the C-terminal fragment of ubiquitin (Cub) tagged with GST-6xHA (Cub-GST) (Fig. 4A). The resulting *GAP1-Cub-GST* was placed under the control of the *GAL1*-promoter. When co-expressed, productive interactions between Gap1 and Shr3 enable the split NubA and Cub domains to assemble a functional ubiquitin moiety that is recognized by ubiquitin-specific proteases, resulting in the release of the GST-6xHA reporter (Fig. 4B). The functional attributes of the NubA fusion constructs were tested by their ability to complement *shr3*Δ (Fig. 4C, dilutions 3-5); strains carrying *SHR3-NubA* or *SHR3*Δ*94-NubA* grew as well as *SHR3* without NubA (Fig. 4C, compare dilution 2 with 3 and 5), whereas the *shr3-35-NubA* allele did not (dilution 4). The *GAP1-Cub-GST* allele encodes a functional Gap1 protein that facilitates citrulline uptake as well as wildtype Gap1 (Fig. 4C, compare dilution 7 with 8). The functionality of Gap1-Cub-GST is dependent on its ability to exit the ER; a construct lacking the ER exit motif in the hydrophilic C-terminal domain of Gap1 is not functional (Fig. 4C dilution 9), presumably due its retention in the ER.

**Figure 4.**
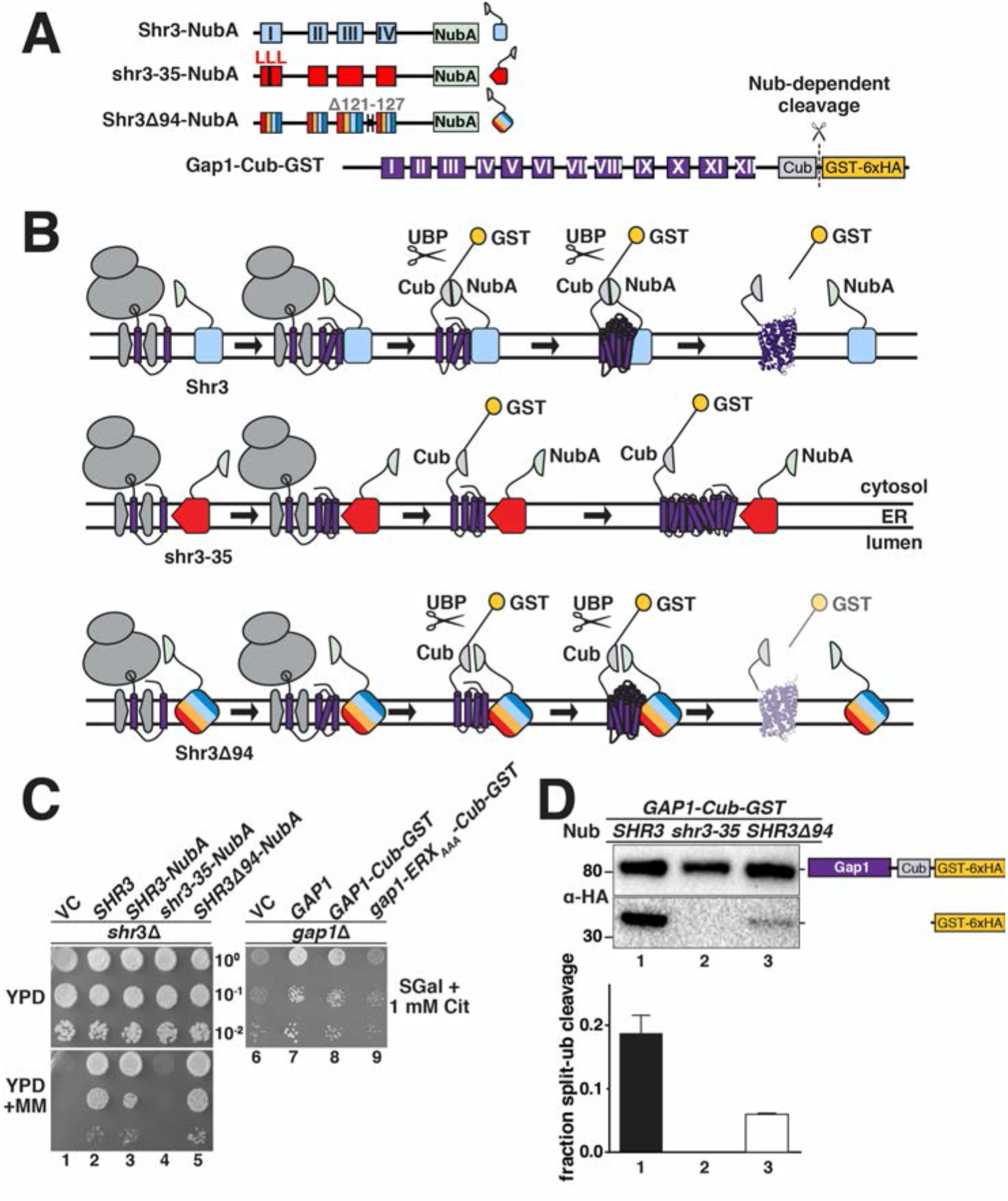
Assessing Shr3-Gap1 interactions using split ubiquitin. (A) Schematic diagram of the split ubiquitin Shr3-NubA, shr3-35-NubA, Shr3Δ94-NubA and Gap1-Cub-GST constructs. (B) Overview of the split-ubiquitin assay and expected outcomes. (C) Left panels: serial dilutions of cell suspensions from strain JKY2 (*shr3*Δ) carrying pRS316 (VC), pPL210 (*SHR3*), pPL1262 (*SHR3-NubA*), pAR67 (*shr3-35-NubA*) or pAR76 (*SHR3*Δ*94-NubA*) spotted on YPD and YPD+MM plates. Plates were incubated at 30 °C for 2 d and photographed. Right panel: serial dilutions of cell suspensions from strain FGY15 (*gap1*Δ) carrying pRS317 (VC), pJK92 (*GAP1*), pPL1257 (*GAP1-Cub-GST*) or pIM28 (*gap1-ERXAAA-Cub-GST*) were spotted on minimal medium with 2 % galactose as carbon source and 1 mM L-citrulline as sole nitrogen source. Plates were incubated for 7 d and photographed. (D) Strain FGY135 (*gap1*Δ *shr3*Δ) expressing *SHR3-NubA* (pPL1262), *shr3-35-NubA* (pAR67) or *SHR3*Δ*94-NubA* (pAR76) and carrying pPL1257 (*GAP1-Cub-GST*) were induced with 2% galactose for 1 h. Proteins extracts were prepared, separated by SDS-PAGE and analyzed by immunoblotting using α-HA antibody. The signal intensities of the immunoreactive forms of full-length and cleaved Gap1 were quantified. The fraction of split-ubiquitin cleavage was determined; the mean values plotted with error bars showing standard deviation (n=3).

To test *in vivo* interactions, we analyzed protein extracts from strain FGY135 (*shr3*Δ *gap1*Δ) carrying plasmids *GAP1-Cub-GST* and *SHR3-NubA, shr3-35-NubA* or *SHR3*Δ*94-NubA* using anti-HA immunoblot analysis. In cells expressing *SHR3-NubA* or *SHR3*Δ*94-NubA* and *GAP1-Cub-GST-6xHA*, two bands were detected, corresponding to full-length Gap1-Cub-GST-6xHA and the cleaved GST-6xHA (Fig. 4D, lane 1). We calculated the fraction of split-ubiquitin cleavage in the *SHR3-NubA* strain to be ≈ 20% by dividing the intensities of the cleaved band with the intensities from full-length plus cleaved species; whereas in the *SHR3*Δ*94-NubA* strain the cleavage was 5% (Fig. 4D); By contrast, only a single band, full-length Gap1-Cub-GST-6xHA, was detected in extracts from the strain expressing the non-functional *shr3-35-NubA* and *GAP1-Cub-GST-6xHA* constructs (Fig. 4D, lane 2). Although expressed at similar levels, the ER retained gap1-ERXAAA-Cub-GST protein did not exhibit an enhanced propensity to interact with Shr3, suggesting the split-ubiquitin assay primarily monitors transient interactions during AAP biogenesis (Fig. S11).

Based on the success of the split ubiquitin approach to analyze Shr3-Gap1 interactions, we created Cub-GST-6xHA tagged constructs with five additional AAP, i.e., Agp1, Gnp1, Bap2, Can1 and Lyp1, the non-transporting but Shr3-dependent AAP homologue Ssy1 (Klasson *et al*., 1999), and two Shr3-independent sugar transporters (HXT), i.e., the low-affinity glucose transporter Hxt1 and the galactose transporter Gal2 (Fig. 5A). Extracts from strain FGY135 (*shr3*Δ *gap1*Δ) carrying *SHR3-NubA, shr3-35-NubA* or *SHR3*Δ*94-NubA* and a single *AAP-Cub-GST-6xHA* or *HXT-Cub-GST-6xHA* construct were prepared and the levels of the GST-6xHA reporter were determined (Fig. 5B-E). Consistent with the general requirement of Shr3 for AAP folding, robust interactions were detected between Agp1-, Gnp1-, Bap2- and Ssy1-Cub constructs and wildtype Shr3-NubA (Fig. 5B, C). Although Shr3 is required for their functional expression, Can1- and Lyp1-Cub constructs exhibited low levels of reporter cleavage, similar to the Shr3-independent sugar transporters (Fig. 5D, E). These latter observations are consistent with Shr3 functioning in a transient manner, primarily interacting with AAP substrates during early stages of AAP folding. In the context of full-length AAP, the split-ubiquitin signal may reflect weaker post-folding interactions.

**Figure 5.**
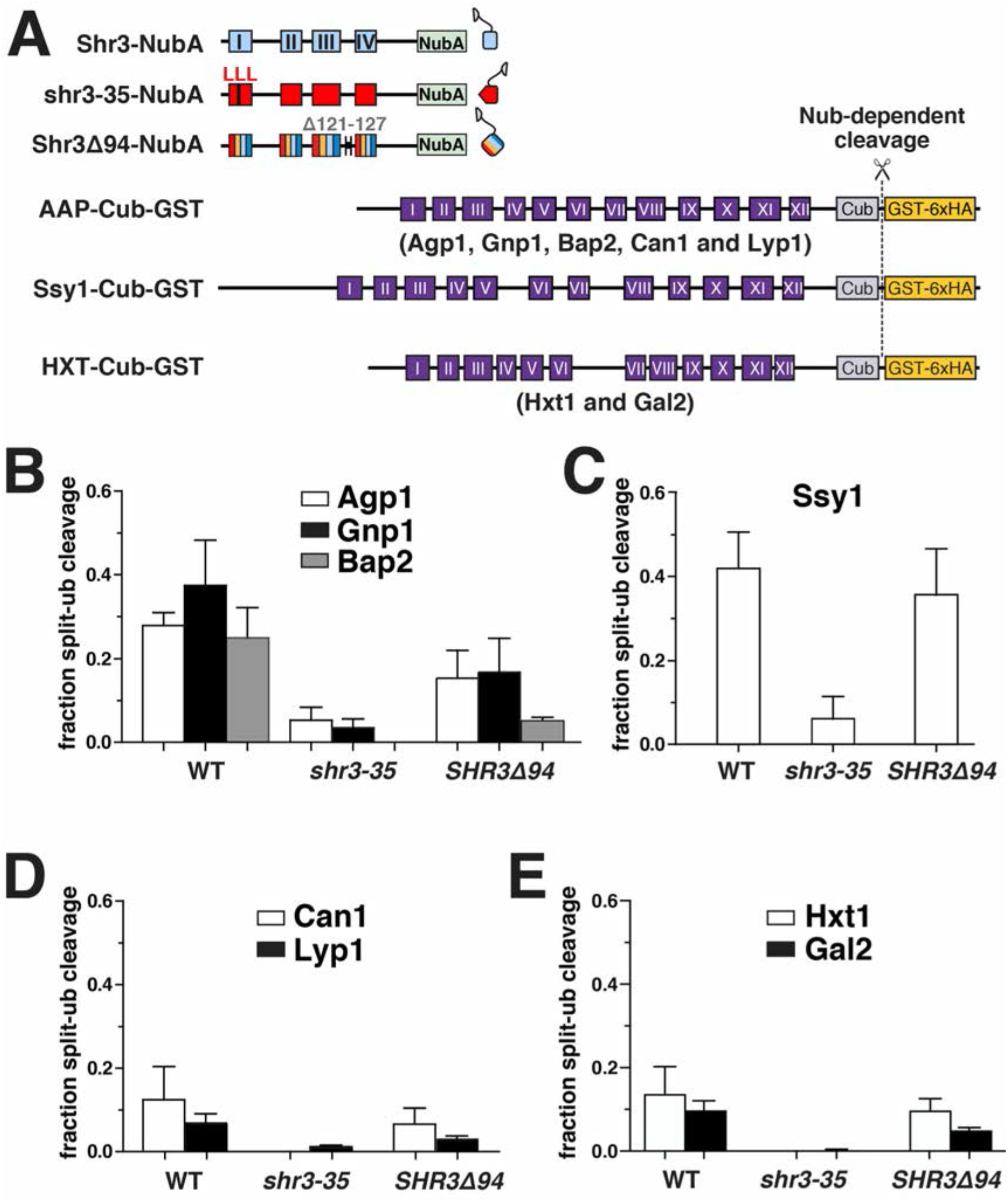
Monitoring Shr3-AAP interactions *in vivo*. (A) Schematic diagram of the split ubiquitin constructs used to evaluate Shr3-AAP and Shr3-HXT interactions. (B) Shr3-Agp1, Shr3-Gnp1 and Shr3-Bap2 interactions. (C) Shr3-Ssy1 interactions. (D) Shr3-Can1 and Shr3-Lyp1 interactions. (E) Shr3-Hxt1 and Shr3-Gal2 interactions. Strain FGY135 (*gap1*Δ *shr3*Δ) expressing *SHR3-NubA* (pPL1262), *shr3-35-NubA* (pAR67) or *SHR3*Δ*94-NubA* (pAR76) and carrying (B) pIM6 (*AGP1-Cub-GST*), pIM17 (*GNP1-Cub-GST*), or pIM7 (*BAP2-Cub-GST*) or (C) pIM19 (*SSY1-Cub-GST*) or (D) pIM8 (*CAN1-Cub-GST*) or pIM18 (*LYP1-Cub-GST*), or (E) pIM32 (*HXT1-Cub-GST*) or pIM33 (*GAL2-Cub-GST*) were induced with 2% galactose for 1 h. Proteins extracts were prepared, separated by SDS-PAGE and analyzed by immunoblotting using α-HA antibody. The signal intensities of the immunoreactive forms of full-length and cleaved Agp1, Gnp1, Bap2, Ssy1, Can1, Lyp1, Hxt1 and Gal2 constructs were quantified. The fraction of split-ubiquitin cleavage was determined; the mean values plotted with error bars showing standard deviation (n=3).

AAP-Cub interactions with shr3-35-NubA were weak or absent, which is precisely aligned with it being non-functional on all selective media tested (Fig. 1, 3). Unexpectedly, the split-Ub interactions with unrelated sugar transporters (HXT) were also weaker compared to the wildtype Shr3-NubA (Fig. 5E). This prompted us to test the trivial explanation that the NubA domain of Shr3-35-NubA is incorrectly oriented. The NubA is correctly oriented to the cytoplasm and thus in a context capable of supporting potential interactions (Supplementary Material, Fig. 12A). Together these findings indicate that the shr3-35 protein, although expressed at similar levels as Shr3, is incapable of engaging in both specific and non-specific secretory substrate interactions.

Interestingly, the Shr3Δ94-NubA interacted with all AAP-Cub constructs, but at significantly reduced levels, and consequently, the data did not explain the enhanced growth conferred by the *SHR3*Δ*94* allele on YPD + MM (Fig. 2 and 3). In marked contrast, Shr3Δ94-NubA interacted with Ssy1-Cub-GST-6xHA at levels comparable to Shr3-NubA (Fig.5 C), which provided the mechanistic explanation for the enhanced growth phenotype. Ssy1 is a unique member of the AAP family that strictly requires Shr3 for folding (Klasson *et al*., 1999) and constitutes the integral membrane component of the PM-localized SPS sensor that induces the expression of AAP genes in response to extracellular amino acids (Ljungdahl and Daignan-Fornier, 2012). Apparently, the Shr3Δ94 mutant retains ability to assist the folding of Ssy1, restoring the transcriptional circuits abrogated by *shr3*Δ. Consistent with this notion, the Shr3Δ94 allele supports Stp1 processing (Fig. 3C). The induced expression of multiple AAP that facilitate branched-chain amino acid uptake correlates well with the growth-based phenotypes. Together, the results indicate that the *in vivo* interactions monitored by the split ubiquitin cleavage provide a nuanced assessment of Shr3 function.

### Shr3 interacts with substrates in a progressive manner

As a proxy to investigate the temporal aspects of Shr3-facilitated AAP folding, we constructed a series of truncated *gap1-Cub-GST, hxt1-Cub-GST* and *gal2-Cub-GST* alleles capable of encoding 2, 4, 6, 8, 10 and 12 MS (Fig. 6A). Strain FGY135 (*shr3*Δ *gap1*Δ) carrying plasmid *SHR3-NubA* and a truncated *gap1-Cub-GST, hxt1-Cub-GST* or *gal2-Cub-GST* allele was employed and potential interactions were monitored by immunoblot. Shr3-NubA did not interact with gap1-2TM, even though the Cub domain is presented in the context of proper membrane topology oriented towards the cytoplasm (Supplementary Material, Fig. S12 B). The presence of two additional MS of Gap1 (gap1-4TM) supported an interaction with Shr3 (Fig. 6 B left panel, C in black). The intensity of the interactions increased and eventually plateaued in the strains carrying the *gap1-6TM/-8TM/-10TM* alleles, respectively (Fig. 6 B left panel, C in black). Strikingly, the gap1-12TM construct interacted only weakly with the functional Shr3-NubA (Fig. 6 B left panel, C in black). In marked contrast to the interaction pattern of the gap1-Cub-GST truncations with Shr3-NubA, we could not detect the GST-6xHA reporter in extracts from the strain expressing *SHR3-NubA* and any of the truncated *hxt1-Cub-GST* or *gal2-Cub-GST* alleles (Fig. 6B, center and left panel, C in grey and white). These findings support the notion that our split ubiquitin approach is suitable to monitor specific Shr3-AAP interactions. As a critical test, we examined if robust interactions could be detected between Shr3-NubA and two truncated Can1 constructs with eight and ten MS (Fig. S13). The rational being that growth-based and biochemical assays have clearly defined Can1 as a *bona fide* substrate of Shr3. However, interactions between full-length Can1 and Shr3 are weak, similar to that corresponding to the non-Shr3 substrates Hxt1 and Gal2 (Fig. 5D and E). We posited that if truncations of AAP are indeed proxies of translation intermediates, then truncations of Can1 would readily interact with Shr3. Interactions between the can1-8TM and -10TM Cub constructs with Shr3-NubA were readily detected (Fig. S13 B, C), findings clearly consistent with Shr3 functioning at early stages of AAP biogenesis.

**Figure 6.**
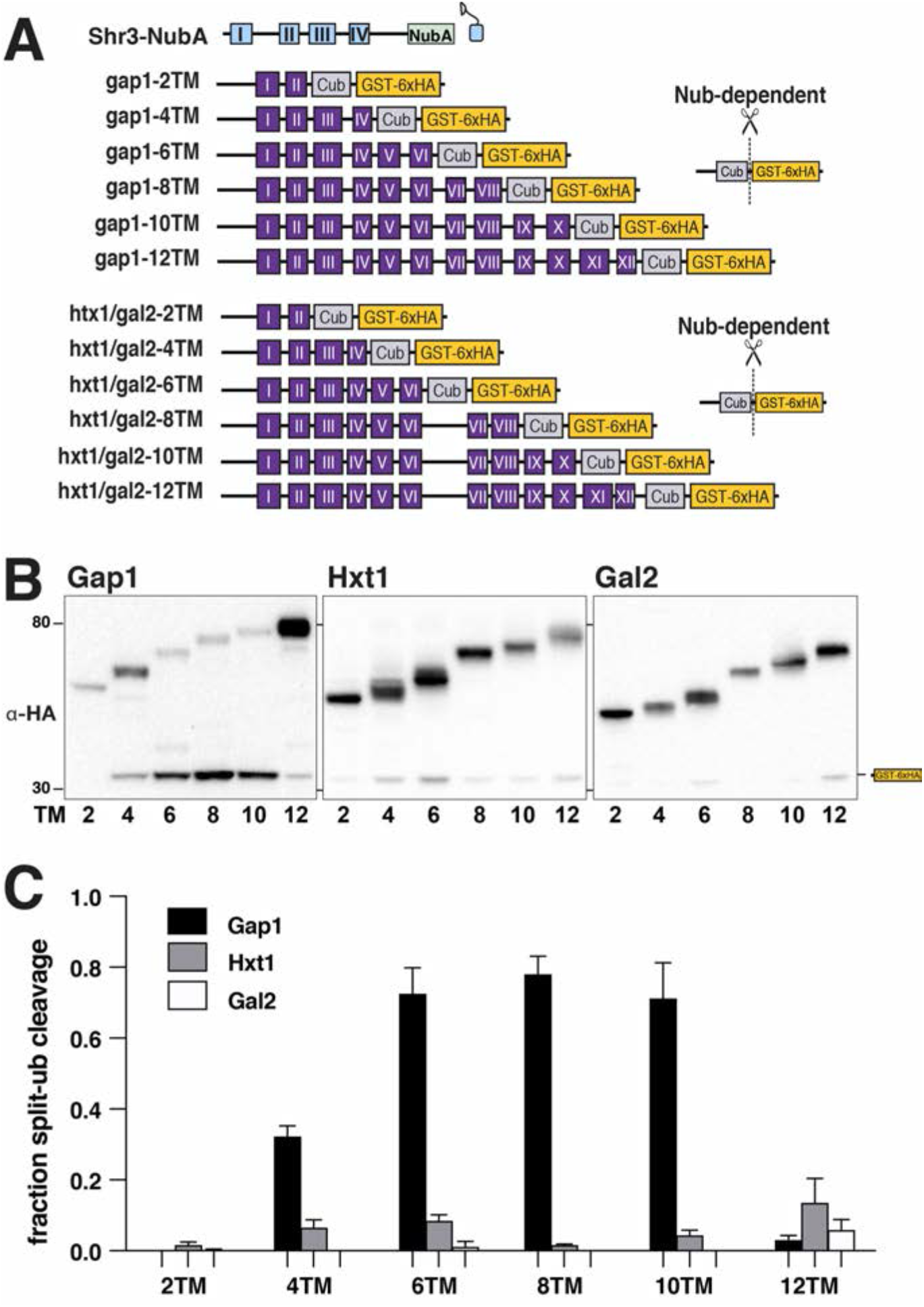
Progressivity of Shr3-Gap1 chaperone-substrate interactions. (A) Schematic diagram of split ubiquitin constructs including the gap1-Cub-GST, hxt1-Cub-GST and gal2-Cub-GST truncation constructs. (B) Strain FGY135 (*gap1*Δ *shr3*Δ) carrying pPL1262 (*SHR3-NubA*) and pIM1 (*gap1-2TM-Cub-GST*), pIM2 (*gap1-4TM-Cub-GST*), pIM3 (*gap1-6TM-Cub-GST*), pIM4 (*gap1-8TM-Cub-GST*), pIM5 (*gap1-10TM-Cub-GST*) or pIM16 (*gap1-12TM-Cub-GST*) (left panel), or pIM34 (*hxt1-2TM-Cub-GST*), pIM35 (*hxt1-4TM-Cub-GST*), pIM36 (*hxt1-6TM-Cub-GST*), pIM37 (*hxt1-8TM-Cub-GST*), pIM38 (*hxt1-10TM-Cub-GST*) or pIM39 (*hxt1-12TM-Cub-GST*) (center panel) or pIM40 (*gal2-2TM-Cub-GST*), pIM41 (*gal2-4TM-Cub-GST*), pIM42 (*gal2-6TM-Cub-GST*), pIM43 (*gal2-8TM-Cub-GST*), pIM44 (*gal2-10TM-Cub-GST*) or pIM45 (*gal2-12TM-Cub-GST*) (right panel) were induced with 2% galactose for 1 h. Extracts were prepared, separated by SDS-PAGE and analyzed by immunoblotting using α-HA antibody. (C) The signal intensities of the immunoreactive forms of uncleaved Cub constructs and cleaved interaction marker (GST-6xHA) were quantified; the mean values of the fraction of split ubiquitin cleavage is plotted with error bars showing standard deviation (n=3).

To more fully understand Shr3-substrate interactions, we created a series of truncated Agp1 and Ssy1 split-ubiquitin constructs (Fig. 7A). Interestingly, in contrast to gap1-2TM, the agp1-2TM construct clearly interacted with Shr3-NubA (Fig. 7A; black bars). Aside from this difference, the pattern of interactions with the remaining Agp1 constructs was strikingly similar to that observed with Gap1 truncations; the intensity of the GST-6xHA reporter increased successively as the number of MS increased from 4 to 10, and greatly reduced when all 12 TM were present (Fig. 7A, black bars). The interactions of the Shr3Δ94-NubA with the agp1-Cub constructs followed a similar pattern to that of the wildtype Shr3-NubA, albeit of lower intensity (Fig. 7A, white bars). Notedly, the interaction pattern with the Ssy1 constructs was quite different (Fig. 7B). Interestingly, as did agp1-2TM, the ssy1-2TM interacted with Shr3-NubA, strongly suggesting that Shr3 engages early during the biogenesis of Ssy1. The ssy1-4TM and -6TM constructs exhibited weaker interactions, however, the ssy1-8TM, -10TM and -12TM constructs exhibited robust interactions. As anticipated from growth-based assays, Shr3Δ94-NubA exhibited an interaction pattern very similar to the wildtype Shr3-NubA construct.

**Figure 7.**
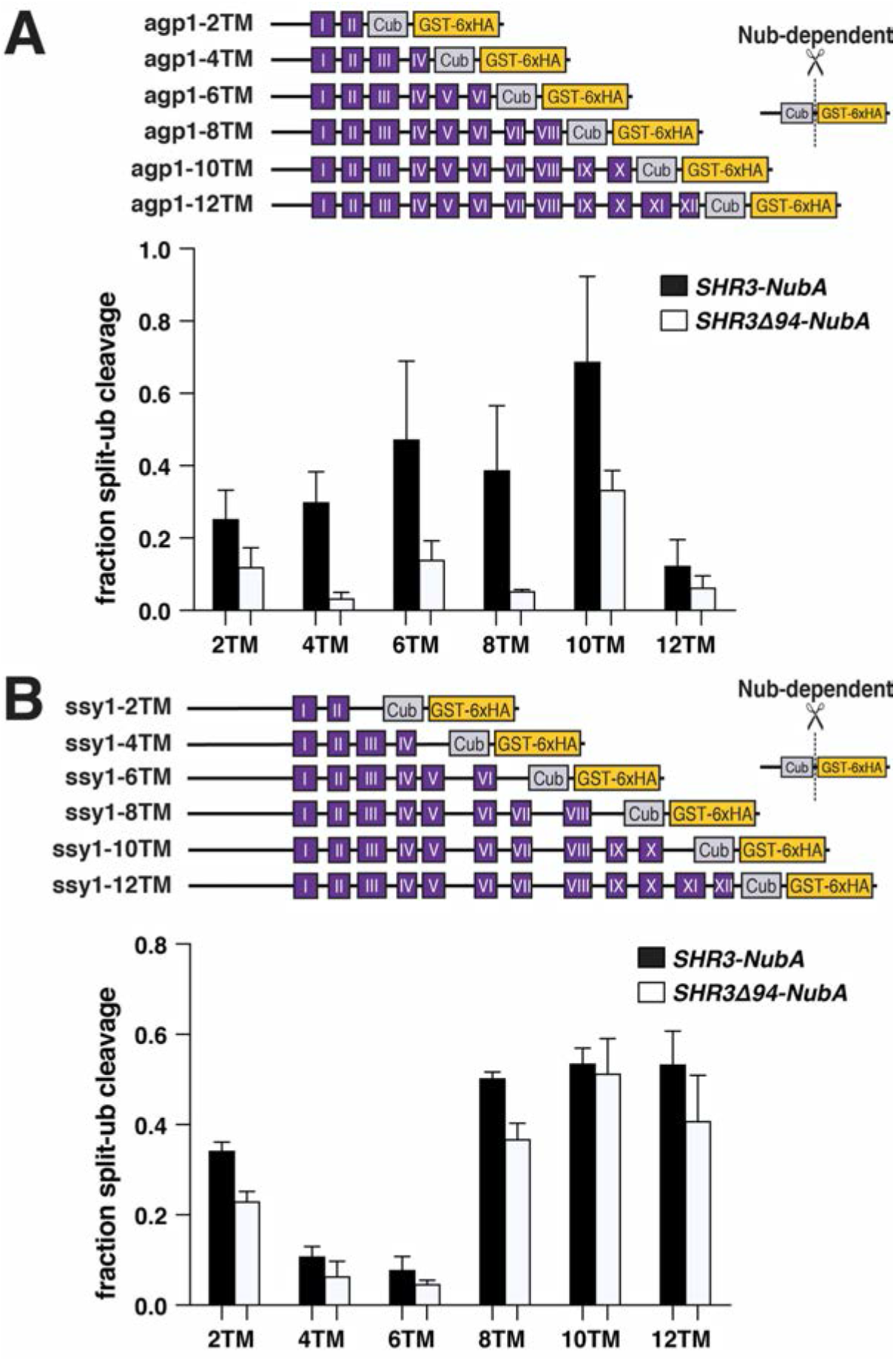
Progressive Shr3-Agp1 and Shr3-Ssy1 chaperone-substrate interactions. (A) Schematic diagram of agp1-Cub-GST truncation constructs. Strain FGY135 (*gap1*Δ *shr3*Δ) expressing *SHR3-NubA* (pPL1262) or *SHR3*Δ*94-NubA* (pAR76) and carrying pIM9 (*agp1-2TM-Cub-GST*), pIM10 (*agp1-4TM-Cub-GST*), pIM11 (*agp1-6TM-Cub-GST*), pIM12 (*agp1-8TM-Cub-GST*), pIM13 (*agp1-10TM-Cub-GST*) or pIM26 (*agp1-12TM-Cub-GST*) were induced with 2% galactose for 1 h. Extracts were prepared, separated by SDS-PAGE and analyzed by immunoblotting using α-HA antibody. The signal intensities of the immunoreactive forms of uncleaved Cub constructs and cleaved interaction marker (GST-6xHA) were quantified; the mean values of the fraction of split ubiquitin cleavage is plotted with error bars showing standard deviation (n=3). (B) Schematic diagram of ssy1-Cub-GST truncation constructs. Strain FGY135 (*gap1*Δ *shr3*Δ) expressing *SHR3-NubA* (pPL1262) or *SHR3*Δ*94-NubA* (pAR76) and carrying pIM20 (*ssy1-2TM-Cub-GST*), pIM21 (*ssy1-4TM-Cub-GST*), pIM22 (*ssy1-6TM-Cub-GST*), pIM23 (*ssy1-8TM-Cub-GST*), pIM24 (*ssy1-10TM-Cub-GST*) or pIM25 (*ssy1-12TM-Cub-GST*) were induced with 2% galactose for 1 h. Extracts were prepared, and analyzed as in (A) and the mean values of the fraction of split ubiquitin cleavage is plotted with error bars showing standard deviation (n=3).

**Figure 8.**
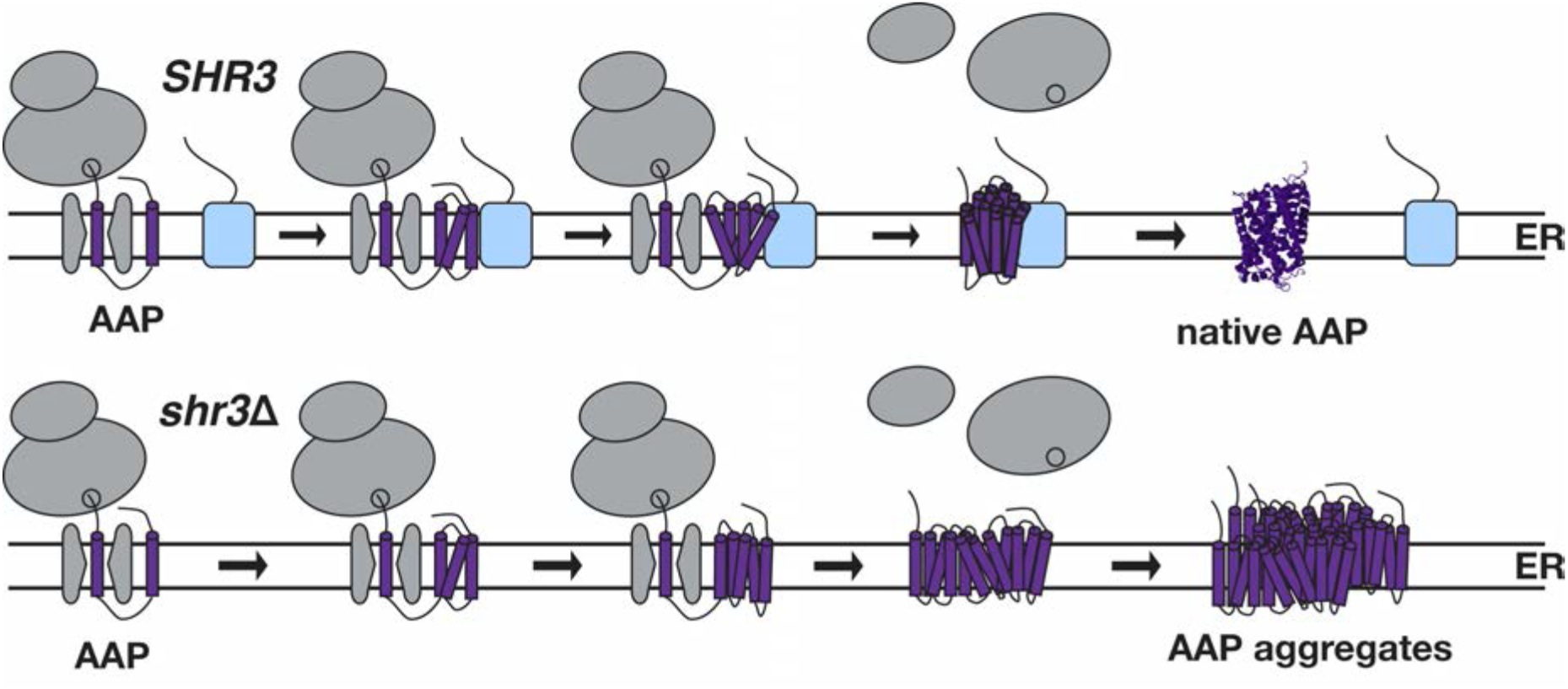
Model of Shr3 facilitated AAP folding. Shr3 interacts transiently with AAP as they are co-translationally inserted into the ER membrane. Interactions start early, when 2-4MS have partitioned into the lipid bilayer, and continue until all MS are inserted. When AAP have fully integrated into the membrane and attain native conformations, the interactions with Shr3 diminish. The co-translational Shr3 function is specifically required for AAP folding and ER-exit. The chaperone activity depends on relatively few residues of the Shr3 sequence, suggesting that it functions as folding template. In the absence of Shr3, AAP are specifically retained in the ER and form high-molecular weight aggregates that are recognized as ERAD substrates.

## Discussion

The biogenesis of AAP can be divided into three interconnected but discrete functional steps: 1) co-translational partitioning and integration into the ER membrane; 2) folding into native structures; and 3) packaging into ER-derived COPII-coated transport vesicles. Shr3 is not essential for integration (Gilstring and Ljungdahl, 2000), but is required for folding and packaging into COPII-coated vesicles (Gilstring *et al*., 1999; Kuehn *et al*., 1998; Kuehn *et al*., 1996); Kota and Ljungdahl, 2005;(Kota *et al*., 2007; Kota and Ljungdahl, 2005). Here, our studies were aimed at further elucidating the role of Shr3 in facilitating AAP folding, i.e. its membrane-localized chaperone (MLC) function.

Saturation scanning mutagenesis of the N-terminal membrane domain of Shr3 allowed us to define amino acid residues that are critical for the recognition of AAP as folding substrates (Supplementary Material Fig. S1-S10; Fig. 3A). Strikingly, mutations affecting a few amino acids residues at three discrete sites resulted in a complete loss of function, suggesting that Shr3 generally recognizes its folding substrates independently of sequence-specific interactions, but rather based on the presence of structural determinants shared by the AAP. The critical residues identified in MSI and MSIV shared the common feature of being polar (Fig. 1C). Threonine19 is conserved among closely and more distant fungal species (Fig. 3B), and it is of interest to note that one of the original spontaneous mutations that led to the identification of *SHR3* is a T19R mutation (Ljungdahl *et al*., 1992). Strikingly, two mutant proteins carrying 10 consecutive leucine residues (aa 62-71, Shr3-57, MSII) or 9 (aa 145-153, Shr3-79, MSIV) support functional expression of AAP in manner indistinguishable from wildtype Shr3, and the Shr3-58 mutant with 13 consecutive leucine residues (aa 69-81, MSII) functions well for all AAP except Gap1 (Fig. 3A, Fig. S6). These findings suggest that hydrophobic interactions between MS of Shr3 and AAP can develop even at the expense of larger sections of specific Shr3 sequence, presumably provided that its overall membrane structure is retained. The data are consistent with Shr3 acting as a scaffold for AAP folding, a function that primarily depends on hydrophobic interactions but with the capacity to shield energetically unfavorable polar residues of AAP MS. Importantly, exposed polar residues may be recognized by a hydrophilic pocket identified in the structure of the ERAD-associated E3 ubiquitin ligase Hrd1 (Schoebel et al., 2017), which participates in the degradation of misfolded AAP in cells lacking Shr3 (Kota *et al*., 2007).

The finding that one of the three loss-of-function mutations disrupting Shr3 function resides in the lumen-oriented loop L1 (Fig. 1) suggests that the role of Shr3 as a MLC is not restricted to MS-mediated hydrophobic interactions. Apparently, Shr3 facilitates the folding of AAP in a manner dependent on extramembrane lumen-oriented sequences. Consistent with this, all internal deletions in L1 of Shr3 strongly affected its function (Fig. 2). Interestingly, L1 contains a predicted α-helical structure with amphipathic characteristics (aa 44-57) that is disrupted in each of the internal deletion mutant proteins.

By contrast to mutations affecting L1, the deletion affecting L3 (*SHR3*Δ*94*) exhibited a variable effect on amino acid uptake (Fig. 2), a phenotype that we could trace to differential interactions with distinct AAP substrates. Consequently, the data implicate L3 as an important determinant that influences substrate interactions. This region appears to be critical for interactions with Gap1 and Can1, but less important for interactions with Agp1/Gnp1, and clearly dispensable for interactions with Ssy1 (Fig. 2, Fig. 4, Fig. 5). Consistently, some mutations in *SHR3* result in WT+ phenotypes, i.e., exhibiting more robust growth than wildtype (Fig. 3A). This latter gain-of-function phenotype is presumably due to proper Ssy1 folding; cells that carry these mutations remain capable of inducing AAP gene expression in a manner that is augmented by the derepression of nitrogen regulation (Ljungdahl and Daignan-Fornier, 2012). Thus, under the specific growth conditions used, cells carrying L3 mutations express enhanced levels of AAP leading to WT+ growth phenotype.

The observation that Ssy1, the only non-transporting AAP, exhibits a lax requirement for L3 to fold, suggests that transporting AAP may have an enhanced requirement for the chaperone function of Shr3. This notion is consistent with recent evidence showing that extracellular loops of AAP are not merely MS-connecting sequences but rather have important roles affecting intracellular trafficking and transport function (van’t Klooster et al., 2020). Some of the hydrophilic loops of AAP are of considerable length, especially extracellular loops 3 and 4 connecting MS V-VI and VII-VIII, respectively. Secondary structure predictions suggest the extracellular loop 4 of the lysine permease Lyp1 (van’t Klooster *et al*., 2020) and Gap1 (Ghaddar et al., 2014a) possesses α-helical regions that appear to influence the substrate specificity of amino acid transport (Risinger et al., 2006). Consistent with a requirement for the lumen-oriented loops of Shr3, the loop regions of AAP that face the extracellular milieu are lumen-oriented during biogenesis. Interactions between the lumen-oriented loops of Shr3 may prevent precocious folding of the extracellular loops, maintaining them in a more flexible state required for subsequent folding events, e.g., involving MS that fold in a context with more distal C-terminal MS. In analogy, proper trafficking and functional expression of the closest AAP homologues in mammals, the L-type amino acid transporters (LAT) (SLC7 family) depend on extramembrane region-mediated recognition by the 4F2hc, or rBAT, members of the SLC3 protein family (Fotiadis et al., 2013). The extracellular orientation of the interacting regions upon plasma-membrane localization of the SLC7-SLC3 holo-LAT transporters in in line with the concept of extramembrane regions containing motifs for substrate recognition and specificity exhibited by amino acid transporter biogenesis factors (Rosell et al., 2014).

Although there is no crystal structure of an AAP, Bap2 (Usami et al., 2014), Can1 (Ghaddar *et al*., 2014a), Gap1 (Ghaddar et al., 2014b) and Tat2 (Kanda and Abe, 2013) have been successfully modeled onto the *E. coli* arginine/agmatine antiporter AdiC (Gao et al., 2010). AdiC has 12 MS and belongs to the Amino Acid-Polyamine-Organocation (APC) super-family of transporters. The 12 MS are arranged in a 5+5 inverted repeat fold that form the transporter core, with MS XI and XII appearing to hold the two halves together. MS I, III, VI, VIII and X shape the binding pocket. To directly probe the *in vivo* interactions between Shr3 and AAP we adapted and applied a split-ubiquitin approach that is independent of a transcriptional readout.

In the context of full-length AAP, we found that the split-ubiquitin signal was relatively weak, specifically in comparison to the signals with truncated substrates. Also, we detected low levels of interactions with full length non-Shr3 substrates that do not rely on Shr3 chaperone function for folding, indicating that the split-ubiquitin approach is sensitive and accurately reflects biologically relevant interactions. Perhaps, Shr3 can interact with many secretory substrates but some interactions become stronger and specific when structural characteristics distinct from the general ones are involved. In support of this notion, the pattern of interactions between Shr3 and truncated Gap1, Agp1, Can1 and Ssy1 is striking, particularly in contrast to truncated HXT (Hxt1 and Gal2) exhibiting essentially no interactions (Fig. 6, 7, and S13). The specific interactions with AAP were detected when only the first 2 to 4 MS were present. As more MS were added the interactions increased and plateaued until MS XI and XII were present, at which point the interactions significantly lessen. By contrast, full-length (Fig. 5) and 12MS truncated (Fig. 7) forms of Ssy1 exhibited more persistent interactions with Shr3 and Shr3Δ94, perhaps attributable to Ssy1 being substantially larger than transporting AAP.

Together, the data we acquired in this study provide further support to our previously described model (Kota *et al*., 2007; Kota and Ljungdahl, 2005) whereby Shr3 transiently interacts with AAP early as their MS partition into the ER membrane, acting as an assembly site for MS helices. This activity is required to prevent AAP translation intermediates from engaging in nonproductive interactions, shielding polar residues, until all MS are available and presented in a context of the long-range intramolecular interactions inherent to the native 3D-structure of AAP (Fig.8). Our model for the Shr3 chaperone function in the biogenesis of AAP is analogous to that of the bacterial insertase/chaperone YidC (Beck et al., 2001; (Dalbey and Kuhn, 2014) acting as an assembly site for alpha helices in the folding of LacY whereby hydrophobic interactions mediate shielding of LacY to provide a protective chamber that reduces energetically unfavorable contacts in the non-native structure during translation (Nagamori et al., 2004; Serdiuk et al., 2016; Serdiuk et al., 2019; Wagner et al., 2008; Zhu et al., 2013). Similar transient MLC-substrate interactions have been reported in the case of the mammalian PAT intramembrane chaperone complex, comprised of four MS, three contributed by Asterix and one by CCDC47 (Chitwood and Hegde, 2020). Although in these studies an *in vitro* translation system was employed, in contrast to our *in vivo* split-ubiquitin approach, the PAT complex interaction was found to be selective for truncated, immature β1-adrenergic receptor constructs compared to the full-length substrate, which presumably is capable of helix packing and polar residue shielding.

The chaperone-like capacity of the conserved eukaryotic ER membrane protein complex (EMC) insertase has recently been explored (Bai et al., 2020; O’Donnell et al., 2020; Pleiner et al., 2020; Volkmar and Christianson, 2020). The results point to EMC acting in close proximity with nascent polytopic membrane proteins typically enriched for MS containing polar or charged residues, shielding them from degradation during folding (Miller-Vedam *et al*., 2020; Shurtleff *et al*., 2018). Importantly, the EMC can associate with a number of ER-integral substrate-specific chaperones such as Sop4 (Luo *et al*., 2002), Gsf2 (Kota and Ljungdahl, 2005; Sherwood and Carlson, 1999) and Ilm1 (Shurtleff *et al*., 2018). Shr3, although an abundant ER membrane protein, has not been identified as an interacting partner of EMC components (Shurtleff *et al*., 2018). Potentially, Shr3 and the EMC act in proximity to distinct ribosome populations, functioning in a parallel manner, enabling the Sec61 translocon to pair with diverse and distinct sets of partners, and thereby facilitate the efficient biogenesis of more challenging versus canonical substrates of the secretory pathway (O’Keefe and High, 2020).

In the case of Shr3, the early interactions with nascent AAP being inserted in the lipid bilayer together with the ability of Shr3 to interact with COPII components via its C-terminal cytoplasmic tail (Gilstring *et al*., 1999), converge to function as a nexus between AAP folding and packaging into COPII-coated vesicles. The potential network of dynamic interactions in the ER remains to be explored for an integral substrate-specific chaperone as well as what structural determinants in the substrates dictate a remarkable degree of substrate specificity. To this end, the broad collection of mutations that we have acquired in combination to our extended growth-based phenotype analysis and in vivo interaction studies lay the ground for future structural studies that are required to gain insights into the mechanistic details that underlie this substrate-specific MLC function.

## Materials and methods

### Yeast strains and plasmids

Yeast strains and plasmid used are listed in Supplementary Material Tables S1 and S2, respectively.

### Media

Standard media, YPD (yeast extract, peptone, dextrose), SD (synthetic defined with ammonium as nitrogen source and glucose as carbon source) were prepared as previously described (Burke et al., 2000). Ammonia-based synthetic complete dextrose (SC) drop-out medium, were prepared as described (Andréasson and Ljungdahl, 2002) and SAD (synthetic minimal dextrose, with allantoin as sole nitrogen source) was prepared as previously described. Media were made solid with 2% (wt/vol) bacto Agar (Difco), 2% (wt/vol) washed bacto Agar (Difco) or 2% (wt/vol) washed pure Agar where indicated. Sensitivity to 200 µg/ml MM (2-{[({[(4-methoxy-6-methyl)-1,3,5-triazin-2-yl]-amino}carbonyl) amino-]-sulfonyl}-benzoic acid) was tested on YPD as described previously (Jørgensen *et al*., 1998). Sensitivity to 1 mM AzC (azetidine-2-carboxylate), 10 µg/ml DL-ethionine, 50 µg/ml p-Fluoro-DL-phenylalanine and 1 µg/ml L-canavanine was tested on SD. Sensitivity to 0,5% (wt/vol) D-histidine was tested on SAD media made solid with washed pure Agar. Cells were grown over night in SC-uracil medium, cells were then resuspended in water to OD=1, 10-fold dilutions were prepared in water and then spotted on the indicated medium. Plates were then incubated at 30°C for 2–3 d and photographed. Gap1-dependent citrulline uptake was monitored on minimal medium containing 2 % galactose as carbon source, 1 mM L-citrulline as sole nitrogen source and uracil. Media were made solid with washed bacto Agar. Plates were incubated at 30 °C for 7 d and photographed.

### Immunoblot analysis

Whole-cell extracts were prepared under denaturing conditions using NaOH and trichloroacetic acid as described previously (Silve et al., 1991). Proteins were separated using SDS-PAGE and blotted onto Amersham Protran 0.45 µm nitrocellulose membrane (GE Healthcare). The primary antibodies and dilutions were, mouse anti-Dpm1 5C5A7 (Abcam), 1:2500; rat anti-HA-HRP 3F10 (Roche Applied Science), 1:2500-1:5000; mouse anti-Pgk1 22C5D8 (Thermo Fisher Scientific), 1:10000 and rabbit anti-Shr3, 1:9000. Secondary antibodies and dilutions used were, goat anti-mouse-poly-HRP (Thermo Fisher Scientific), 1:5000 and goat anti-rabbit-poly-HRP (Thermo Fisher Scientific), 1:5000. Immunoreactive bands were visualized by chemiluminescence using (SuperSignal West Dura Extended-Duration Substrate; Thermo Fisher Scientific) as substrate in a ChemiDoc imaging system (Biorad).

### Split-ubiquitin assay

Cells were pre-grown in SD+R (synthetic defined with ammonium as nitrogen source and 2 % raffinose and 0.1 % glucose as carbon source) to logarithmic phase. Approximately 10 OD of logarithmically cells were induced in 5 ml of SD+G (synthetic defined with ammonium as nitrogen source and 2 % galactose as carbon source) for 1 hour. Cells were collected and washed once in ddH20. Cells were resuspended in 150 µl lysis buffer (0.8 M sorbitol; 10 mM MOPS, pH 7.2; 2 mM EDTA; 1 mM PMSF; 1X cØmplete, mini, EDTA-free protease inhibitor cocktail, Roche). Cells were lysed by bead beating with 0.5 mm glass beads for 3×20s at 6.5 m/s in a benchtop homogenizer (Fastprep-24, MP Biomedical). The cell lysates were centrifuged at 500g for 10 min and 25 µl of the resulting supernatant was diluted 1:1 with 2x sample buffer. Proteins were separated using SDS-PAGE and blotted onto Amersham Protran 0.45 µm nitrocellulose membrane (GE Healthcare). The primary antibody and dilution used was rat anti-HA-HRP 3F10 (Roche Applied Science), 1:2500-1:5000. Immunoreactive bands were visualized by chemiluminescence using (SuperSignal West Dura Extended-Duration Substrate; Thermo Fisher Scientific) as substrate in a ChemiDoc imaging system (Biorad).

### Protease protection assay

Cells were pre-grown in SD+R (synthetic defined with ammonium as nitrogen source and 2 % raffinose and 0.1 % glucose as carbon source) to logarithmic phase. Approximately 5 OD of logarithmically cells were induced in 5 ml of SD+G (synthetic defined with ammonium as nitrogen source and 2 % galactose as carbon source) for 1 hour. Cells were collected and washed once in ddH20. Cells were resuspended in 150 µl lysis buffer (0.8 M sorbitol; 10 mM MOPS, pH 7.2; 2 mM EDTA; 1 mM PMSF; 1X cØmplete, mini, EDTA-free protease inhibitor cocktail, Roche). Cells were lysed by bead beating with 0.5 mm glass beads for 3×20s at 6.5 m/s in a benchtop homogenizer (Fastprep-24, MP Biomedical). The cell lysates were centrifuged at 500g for 10 min and 100 µl of the resulting supernatant was centrifuged at 100 000 g for 30 minutes. The membrane pellet was resuspended in 50 µl lysis buffer (0.8 M sorbitol; 10 mM MOPS, pH 7.2; 2 mM EDTA; 5 mM CaCl2). The resulting membrane preparations were digested with 20 µg Proteinase K (Thermo Fisher Scientific) on ice with 0.2 % NP-40 as indicated. Time points where taken at 0 and 2 h. Proteins were precipitated using trichloroacetic acid as described previously (Silve *et al*., 1991). Proteins were separated using SDS-PAGE and blotted onto Amersham Protran 0.45 µm nitrocellulose membrane (GE Healthcare). The primary antibodies and dilutions used were, rat anti-HA-HRP 3F10 (Roche Applied Science), 1:2500-1:5000 and rabbit anti-Kar2, 1:5000. Secondary antibody and dilution used was goat anti-rabbit-poly-HRP (Thermo Fisher Scientific), 1:5000. Immunoreactive bands were visualized by chemiluminescence using (SuperSignal West Dura Extended-Duration Substrate; Thermo Fisher Scientific) as substrate in a ChemiDoc imaging system (Biorad).

## Acknowledgements

We thank the members of the Ljungdahl laboratory and Claes Andréasson for constructive comments throughout the course of this work. In particular we acknowledge Nina Horwege, and Carlos Sacristán for early contributions in creating plasmid constructs. This research was supported by funding from Swedish Research Council (P.O.L.), Grant/Award numbers: 2011-5925 and 2015-04202.

## Competing interests

The authors have no conflicts of interest to report.

## Supplementary Material

### Figures

**Fig S1**. Growth-based assessment of Shr3 substrate specificity - I

**Fig S2**. Growth-based assessment of Shr3 substrate specificity - II

**Fig S3**. Growth-based assessment of Shr3 substrate specificity - III

**Fig S4**. Growth-based assessment of Shr3 substrate specificity - IV

**Fig S5**. Growth-based assessment of Shr3 substrate specificity - V

**Fig S6**. Growth-based assessment of Shr3 substrate specificity - VI

**Fig S7**. Growth-based assessment of Shr3 substrate specificity - VII

**Fig S8**. Growth-based assessment of Shr3 substrate specificity - VIII

**Fig S9**. Growth-based assessment of Shr3 substrate specificity - IX

**Fig S10**. Growth-based assessment of Shr3 substrate specificity - X

**Fig S11**. Effect of ER exit motif mutations on Shr3-AAP interactions

**Fig S12**. Protease cleavage assay to assess the topology of shr3-35-NubA and gap1-2TM-Cub-GST constructs

**Fig S13**. Interactions between Shr3-NubA and can1-8TM, -10TM-Cub-GST

### Tables

**Table S1**. Strains

**Table S2**. Plasmids

## Notes

### Competing Interest Statement

The authors have declared no competing interest.

### Summary of Updates

To better gauge the observed protein-protein interactions described, we have carried out several additional and important controls. Shr3 interacts with full-length amino acid permease substrates, but not with two non-substrate sugar transporters Gal2 and Hxt1 (new data, revised Fig. 5). Importantly, we also found that full-length amino acid permeases interact less well than truncated versions of amino acid permeases, e.g., Can1 is a striking example of this (new Fig. S13). The latter results are consistent with the truncated forms of amino acid permeases being proxies of translation intermediates. This notion is strongly supported by the finding that truncated versions of the sugar transporter do not interact with Shr3 (new data, revised Fig. 6), again underscoring the specificity exhibited by Shr3 for its substrates. The data indicating that full-length, presumably folded, amino acid permeases exhibit a lower propensity to interact with Shr3 (revised Fig. 6, 7, S13) are consistent with the transient nature of Shr3-substrate interactions. In summary, the inclusion of new data provides unique mechanistic insights and provide a solid foundation for understanding the known selectively of this paradigm membrane-localized chaperone.

## References

Andréasson, C., and Ljungdahl, P.O. (2002). Receptor-mediated endoproteolytic activation of two transcription factors in yeast. Genes Dev. 16, 3158–3172.

Andréasson, C., Neve, E.P.A., and Ljungdahl, P.O. (2004). Four permeases import proline and the toxic proline analogue azetidine-2-carboxylate into yeast. Yeast 21, 193–199.

Bai, L., You, Q., Feng, X., Kovach, A., and Li, H. (2020). Structure of the ER membrane complex, a transmembrane-domain insertase. Nature 584, 475–478. 10.1038/s41586-020-2389-3.

Chitwood, P.J., and Hegde, R.S. (2020). An intramembrane chaperone complex facilitates membrane protein biogenesis. Nature 584, 630–634. 10.1038/s41586-020-2624-y.

Dalbey, R.E., and Kuhn, A. (2014). How YidC inserts and folds proteins across a membrane. Nat Struct Mol Biol 21, 435–436. 10.1038/nsmb.2823.

Didion, T., Regenberg, B., Jørgensen, M.U., Kielland-Brandt, M.C., and Andersen, H.A. (1998). The permease homologue Ssy1p controls the expression of amino acid and peptide transporter genes in Saccharomyces cerevisiae. Mol. Microbiol. 27, 643–650.

Drozdetskiy, A., Cole, C., Procter, J., and Barton, G.J. (2015). JPred4: a protein secondary structure prediction server. Nucleic Acids Res 43, W389–394. 10.1093/nar/gkv332.

Erpapazoglou, Z., Kafasla, P., and Sophianopoulou, V. (2006). The product of the SHR3 orthologue of Aspergillus nidulans has restricted range of amino acid transporter targets. Fungal Genet Biol 43, 222–233. 10.1016/j.fgb.2005.11.006.

Fotiadis, D., Kanai, Y., and Palacin, M. (2013). The SLC3 and SLC7 families of amino acid transporters. Mol Aspects Med 34, 139–158. 10.1016/j.mam.2012.10.007.

Gao, X., Zhou, L., Jiao, X., Lu, F., Yan, C., Zeng, X., Wang, J., and Shi, Y. (2010). Mechanism of substrate recognition and transport by an amino acid antiporter. Nature 463, 828–832. 10.1038/nature08741.

Ghaddar, K., Krammer, E.M., Mihajlovic, N., Brohee, S., Andre, B., and Prevost, M. (2014a). Converting the yeast arginine can1 permease to a lysine permease. J Biol Chem 289, 7232–7246. 10.1074/jbc.M113.525915.

Ghaddar, K., Merhi, A., Saliba, E., Krammer, E.M., Prevost, M., and Andre, B. (2014b). Substrate-induced ubiquitylation and endocytosis of yeast amino acid permeases. Mol Cell Biol 34, 4447–4463. 10.1128/MCB.00699-14.

Gilstring, C.F., and Ljungdahl, P.O. (2000). A method for determining the in vivo topology of yeast polytopic membrane proteins demonstrates that Gap1p fully integrates into the membrane independently of Shr3p. J. Biol. Chem. 275, 31488–31495.

Gilstring, C.F., Melin-Larsson, M., and Ljungdahl, P.O. (1999). Shr3p mediates specific COPII coatomer-cargo interactions required for the packaging of amino acid permeases into ER-derived transport vesicles. Mol. Biol. Cell 10, 3549–3565.

Grenson, M., Hou, C., and Crabeel, M. (1970). Multiplicity of the amino acid permeases in Saccharomyces cerevisiae. IV. Evidence for a general amino acid permease. J Bacteriol 103, 770–777.

Gresham, D., Usaite, R., Germann, S.M., Lisby, M., Botstein, D., and Regenberg, B. (2010). Adaptation to diverse nitrogen-limited environments by deletion or extrachromosomal element formation of the GAP1 locus. Proc Natl Acad Sci U S A 107, 18551–18556. 10.1073/pnas.1014023107.

Heinrich, S.U., Mothes, W., Brunner, J., and Rapoport, T.A. (2000). The Sec61 complex mediates the integration of a membrane protein by allowing lipid partitioning of the transmembrane domain. Cell 102, 233–244.

Heinrich, S.U., and Rapoport, T.A. (2003). Cooperation of transmembrane segments during the integration of a double-spanning protein into the ER membrane. Embo J 22, 3654–3663.

Hou, B., Lin, P.J., and Johnson, A.E. (2012). Membrane protein TM segments are retained at the translocon during integration until the nascent chain cues FRET-detected release into bulk lipid. Mol Cell 48, 398–408. 10.1016/j.molcel.2012.08.023.

Iraqui, I., Vissers, S., Bernard, F., de Craene, J.O., Boles, E., Urrestarazu, A., and André, B. (1999). Amino acid signaling in Saccharomyces cerevisiae: a permease-like sensor of external amino acids and F-Box protein Grr1p are required for transcriptional induction of the AGP1 gene, which encodes a broad-specificity amino acid permease. Mol. Cell. Biol. 19, 989–1001.

Jack, D.L., Paulsen, I.T., and Saier, M.H. (2000). The amino acid/polyamine/organocation (APC) superfamily of transporters specific for amino acids, polyamines and organocations. Microbiology 146 (Pt 8), 1797–1814. 10.1099/00221287-146-8-1797.

Johnson, A.E., and van Waes, M.A. (1999). The translocon: a dynamic gateway at the ER membrane. Annu Rev Cell Dev Biol 15, 799–842. 10.1146/annurev.cellbio.15.1.799.

Johnsson, N. (2002). A split-ubiquitin-based assay detects the influence of mutations on the conformational stability of the p53 DNA binding domain in vivo. FEBS Lett 531, 259-264. S0014579302035330 [pii].

Johnsson, N., and Varshavsky, A. (1994). Split ubiquitin as a sensor of protein interactions in vivo. Proc Natl Acad Sci U S A 91, 10340–10344.

Jørgensen, M.U., Bruun, M.B., Didion, T., and Kielland-Brandt, M.C. (1998). Mutations in five loci affecting GAP1-independent uptake of neutral amino acids in yeast. Yeast 14, 103–114.

Kanda, N., and Abe, F. (2013). Structural and functional implications of the yeast high-affinity tryptophan permease Tat2. Biochemistry 52, 4296–4307. 10.1021/bi4004638.

Klasson, H., Fink, G.R., and Ljungdahl, P.O. (1999). Ssy1p and Ptr3p are plasma membrane components of a yeast system that senses extracellular amino acids. Mol Cell Biol 19, 5405–5416.

Kota, J., Gilstring, C.F., and Ljungdahl, P.O. (2007). Membrane chaperone Shr3 assists in folding amino acid permeases preventing precocious ERAD. J Cell Biol 176, 617–628. 10.1083/jcb.200612100.

Kota, J., and Ljungdahl, P.O. (2005). Specialized membrane-localized chaperones prevent aggregation of polytopic proteins in the ER. J Cell Biol 168, 79–88.

Kuehn, M.J., Herrmann, J.M., and Schekman, R. (1998). COPII-cargo interactions direct protein sorting into ER-derived transport vesicles. Nature 391, 187–190.

Kuehn, M.J., Schekman, R., and Ljungdahl, P.O. (1996). Amino acid permeases require COPII components and the ER resident membrane protein Shr3p for packaging into transport vesicles in vitro. J. Cell Biol. 135, 585–595.

Lau, W.T., Howson, R.W., Malkus, P., Schekman, R., and O’Shea, E.K. (2000). Pho86p, an endoplasmic reticulum (ER) resident protein in Saccharomyces cerevisiae, is required for ER exit of the high-affinity phosphate transporter Pho84p. Proc. Natl. Acad. Sci. USA 97, 1107–1112.

Ljungdahl, P.O., and Daignan-Fornier, B. (2012). Regulation of amino acid, nucleotide, and phosphate metabolism in Saccharomyces cerevisiae. Genetics 190, 885–929. 10.1534/genetics.111.133306.

Ljungdahl, P.O., Gimeno, C.J., Styles, C.A., and Fink, G.R. (1992). SHR3: A novel component of the secretory pathway specifically required for the localization of amino acid permeases in yeast. Cell 71, 463–478.

Luo, W.J., Gong, X.H., and Chang, A. (2002). An ER membrane protein, Sop4, facilitates ER export of the yeast plasma membrane [H+]ATPase, Pma1. Traffic 3, 730–739. 10.1034/j.1600-0854.2002.31005.x.

Madeira, F., Park, Y.M., Lee, J., Buso, N., Gur, T., Madhusoodanan, N., Basutkar, P., Tivey, A.R.N., Potter, S.C., Finn, R.D., and Lopez, R. (2019). The EMBL-EBI search and sequence analysis tools APIs in 2019. Nucleic Acids Research 47, W636–W641. https://doi.org/10.1093/nar/gkz268.

Malkus, P., Jiang, F., and Schekman, R. (2002). Concentrative sorting of secretory cargo proteins into COPII-coated vesicles. J. Cell Biol. 159, 915–921.

Martínez, P., and Ljungdahl, P.O. (2000). The SHR3 homologue from S. pombe demonstrates a conserved function of ER packaging chaperones. J. Cell Sci. 113, 4351–4362.

Martínez, P., and Ljungdahl, P.O. (2004). An ER packaging chaperone determines the amino acid uptake capacity and virulence of Candida albicans. Mol Microbiol 51, 371–384.

Miller, E., Antonny, B., Hamamoto, S., and Schekman, R. (2002). Cargo selection into COPII vesicles is driven by the Sec24p subunit.EMBO J. 21, 6105–6113.

Miller, E.A., Beilharz, T.H., Malkus, P.N., Lee, M.C., Hamamoto, S., Orci, L., and Schekman, R. (2003). Multiple cargo binding sites on the COPII subunit Sec24p ensure capture of diverse membrane proteins into transport vesicles. Cell 114, 497–509.

Miller-Vedam, L.E., Brauning, B., Popova, K.D., Schirle Oakdale, N.T., Bonnar, J.L., Prabu, J.R., Boydston, E.A., Sevillano, N., Shurtleff, M.J., Stroud, R.M., et al. (2020). Structural and mechanistic basis of the EMC-dependent biogenesis of distinct transmembrane clients. Elife 9. 10.7554/eLife.62611.

Nagamori, S., Smirnova, I.N., and Kaback, H.R. (2004). Role of YidC in folding of polytopic membrane proteins. J. Cell Biol. 165, 53–62.

O’Donnell, J.P., Phillips, B.P., Yagita, Y., Juszkiewicz, S., Wagner, A., Malinverni, D., Keenan, R.J., Miller, E.A., and Hegde, R.S. (2020). The architecture of EMC reveals a path for membrane protein insertion. Elife 9. 10.7554/eLife.57887.

O’Keefe, S., and High, S. (2020). Membrane translocation at the ER: with a little help from my friends. FEBS J. 10.1111/febs.15309.

Ono, B.I., Ishino, Y., and Shinoda, S. (1983). Nonsense mutations in the can1 locus of Saccharomyces cerevisiae. J Bacteriol 154, 1476–1479.

Pleiner, T., Tomaleri, G.P., Januszyk, K., Inglis, A.J., Hazu, M., and Voorhees, R.M. (2020). Structural basis for membrane insertion by the human ER membrane protein complex. Science 369, 433–436. 10.1126/science.abb5008.

Rapoport, T.A., Li, L., and Park, E. (2017). Structural and Mechanistic Insights into Protein Translocation. Annu Rev Cell Dev Biol 33, 369–390. 10.1146/annurev-cellbio-100616-060439.

Risinger, A.L., Cain, N.E., Chen, E.J., and Kaiser, C.A. (2006). Activity-dependent reversible inactivation of the general amino acid permease. Mol Biol Cell 17, 4411–4419. 10.1091/mbc.e06-06-0506.

Rosell, A., Meury, M., Alvarez-Marimon, E., Costa, M., Perez-Cano, L., Zorzano, A., Fernandez-Recio, J., Palacin, M., and Fotiadis, D. (2014). Structural bases for the interaction and stabilization of the human amino acid transporter LAT2 with its ancillary protein 4F2hc. Proc Natl Acad Sci U S A 111, 2966–2971. 10.1073/pnas.1323779111.

Rytka, J. (1975). Positive selection of general amino acid permease mutants in Saccharomyces cerevisiae. J Bacteriol 121, 562–570.

Saier, M.H., Jr. (2000). Families of transmembrane transporters selective for amino acids and their derivatives. Microbiology 146 (Pt 8), 1775–1795. 10.1099/00221287-146-8-1775.

Schoebel, S., Mi, W., Stein, A., Ovchinnikov, S., Pavlovicz, R., DiMaio, F., Baker, D., Chambers, M.G., Su, H., Li, D., et al. (2017). Cryo-EM structure of the protein-conducting ERAD channel Hrd1 in complex with Hrd3. Nature 548, 352–355. 10.1038/nature23314.

Seinen, A.B., and Driessen, A.J.M. (2019). Single-Molecule Studies on the Protein Translocon. Annu Rev Biophys 48, 185–207. 10.1146/annurev-biophys-052118-115352.

Serdiuk, T., Balasubramaniam, D., Sugihara, J., Mari, S.A., Kaback, H.R., and Muller, D.J. (2016). YidC assists the stepwise and stochastic folding of membrane proteins. Nat Chem Biol 12, 911–917. 10.1038/nchembio.2169.

Serdiuk, T., Steudle, A., Mari, S.A., Manioglu, S., Kaback, H.R., Kuhn, A., and Muller, D.J. (2019). Insertion and folding pathways of single membrane proteins guided by translocases and insertases. Sci Adv 5, eaau6824. 10.1126/sciadv.aau6824.

Sherwood, P.W., and Carlson, M. (1999). Efficient export of the glucose transporter Hxt1p from the endoplasmic reticulum requires Gsf2p. Proc. Natl. Acad. Sci. USA 96, 7415–7420.

Shurtleff, M.J., Itzhak, D.N., Hussmann, J.A., Schirle Oakdale, N.T., Costa, E.A., Jonikas, M., Weibezahn, J., Popova, K.D., Jan, C.H., Sinitcyn, P., et al. (2018). The ER membrane protein complex interacts cotranslationally to enable biogenesis of multipass membrane proteins. Elife 7. 10.7554/eLife.37018.

Silve, S., Volland, C., Garnier, C., Jund, R., Chevallier, M.R., and Haguenauer-Tsapis, R. (1991). Membrane insertion of uracil permease, a polytopic yeast plasma membrane protein. Mol. Cell. Biol. 11, 1114–1124.

Usami, Y., Uemura, S., Mochizuki, T., Morita, A., Shishido, F., Inokuchi, J., and Abe, F. (2014). Functional mapping and implications of substrate specificity of the yeast high-affinity leucine permease Bap2. Biochim Biophys Acta 1838, 1719–1729. 10.1016/j.bbamem.2014.03.018.

van’t Klooster, J.S., Bianchi, F., Doorn, R.B., Lorenzon, M., Lusseveld, J.H., Punter, C.M., and Poolman, B. (2020). Extracellular loops matter - subcellular location and function of the lysine transporter Lyp1 from Saccharomyces cerevisiae. FEBS J. 10.1111/febs.15262.

Volkmar, N., and Christianson, J.C. (2020). Squaring the EMC - how promoting membrane protein biogenesis impacts cellular functions and organismal homeostasis. J Cell Sci 133. 10.1242/jcs.243519.

Wagner, S., Pop, O.I., Haan, G.J., Baars, L., Koningstein, G., Klepsch, M.M., Genevaux, P., Luirink, J., and de Gier, J.W. (2008). Biogenesis of MalF and the MalFGK(2) maltose transport complex in Escherichia coli requires YidC. J Biol Chem 283, 17881–17890. 10.1074/jbc.M801481200.

Waterhouse, A.M., Procter, J.B., Martin, D.M., Clamp, M., and Barton, G.J. (2009). Jalview Version 2--a multiple sequence alignment editor and analysis workbench. Bioinformatics 25, 1189–1191. 10.1093/bioinformatics/btp033.

Wong, F.H., Chen, J.S., Reddy, V., Day, J.L., Shlykov, M.A., Wakabayashi, S.T., and Saier, M.H., Jr. (2012). The amino acid-polyamine-organocation superfamily. J Mol Microbiol Biotechnol 22, 105–113. 10.1159/000338542.

Zhu, L., Kaback, H.R., and Dalbey, R.E. (2013). YidC protein, a molecular chaperone for LacY protein folding via the SecYEG protein machinery. J Biol Chem 288, 28180–28194. 10.1074/jbc.M113.491613.

