## Supplemental Material for "The ER membrane chaperone Shr3 acts in a progressive manner to assist the folding of related plasma membrane transport proteins"

### Figures:

**Fig S1.** Growth-based assessment of Shr3 substrate specificity - I

**Fig S2.** Growth-based assessment of Shr3 substrate specificity - II

**Fig S3.** Growth-based assessment of Shr3 substrate specificity - III

**Fig S4.** Growth-based assessment of Shr3 substrate specificity - IV

**Fig S5.** Growth-based assessment of Shr3 substrate specificity - V

**Fig S6.** Growth-based assessment of Shr3 substrate specificity - VI

**Fig S7.** Growth-based assessment of Shr3 substrate specificity - VII

**Fig S8.** Growth-based assessment of Shr3 substrate specificity - VIII

**Fig S9.** Growth-based assessment of Shr3 substrate specificity - IX

**Fig S10.** Growth-based assessment of Shr3 substrate specificity - X

**Fig S11.** Effect of ER exit motif mutations on Shr3-AAP interactions

**Fig S12.** Protease cleavage assay to assess the topology of shr3-35-NubA and gap1-2TM-Cub-GST constructs

**Fig S13.** Interactions between Shr3-NubA and can1-8TM, -10TM-Cub-GST

### Tables:

**Table S1.** Strains

**Table S2.** Plasmids

### References

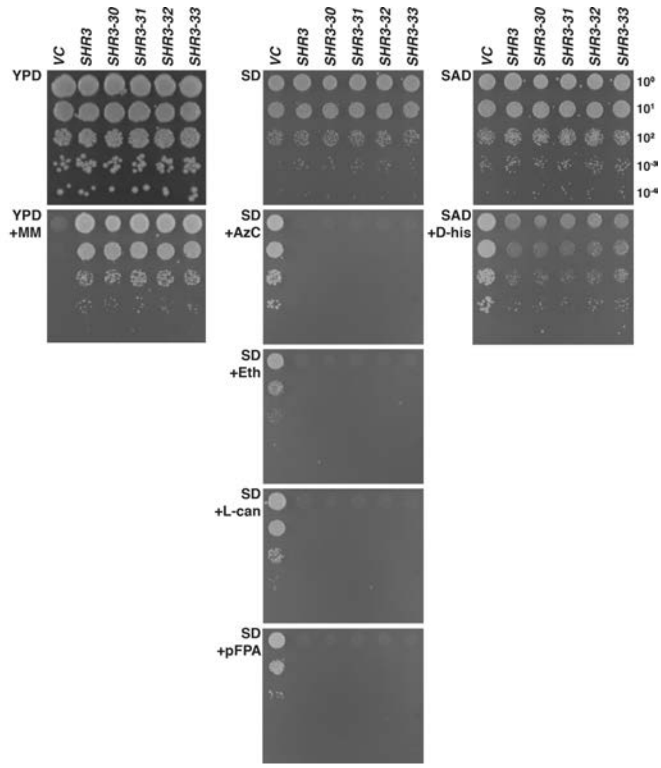


**Figure S1. Growth-based assessment of Shr3 substrate specificity - I**

Growth of serially diluted cell suspensions from strain JKY2 (*shr3*Δ) expressing indicated Shr3-mutant proteins spotted on indicated media. Concentrations of toxic amino analogues and nitrogen sources were as follows: 200 µg/ml metsulfuron-methyl (MM), yeast extract and peptone; 1 mM azetidine-2-carboxylate (AzC), ammonium; 10 µg/ml DL-ethionine (Eth), ammonium; 1 µg/ml L-canavanine (L-can), ammonium; 50 µg/ml p-Fluoro-DL-phenylalanine (pFPA), ammonium; 0.5% w/v D-histidine (D-his), allantoin. Growth was scored after 2 to 3 d of incubation at 30 °C.

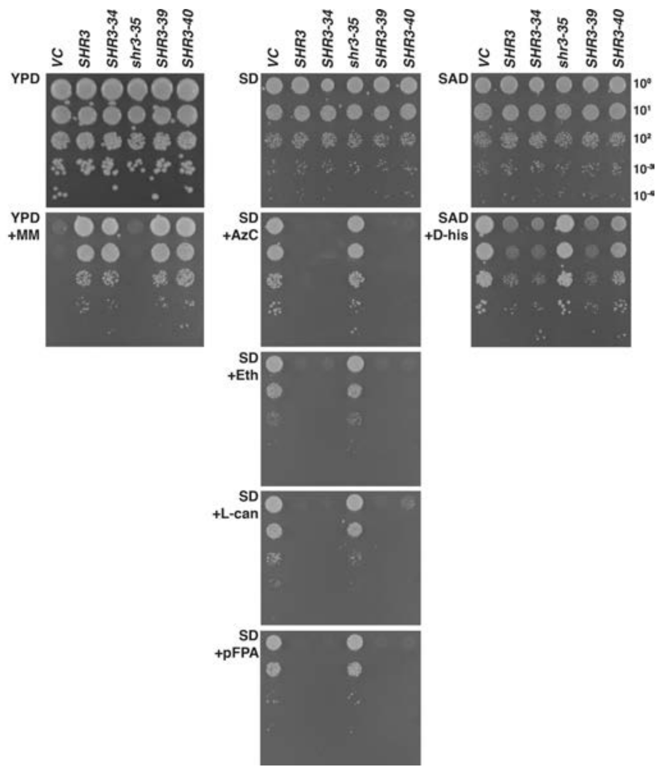


**Figure S2. Growth-based assessment of Shr3 substrate specificity - II**

Growth of serially diluted cell suspensions from strain JKY2 (*shr3*Δ) expressing indicated Shr3-mutant proteins spotted on indicated media. Concentrations of toxic amino analogues and nitrogen sources were as follows: 200 µg/ml metsulfuron-methyl (MM), yeast extract and peptone; 1 mM azetidine-2-carboxylate (AzC), ammonium; 10 µg/ml DL-ethionine (Eth), ammonium; 1 µg/ml L-canavanine (L-can), ammonium; 50 µg/ml p-Fluoro-DL-phenylalanine (pFPA), ammonium; 0.5% w/v D-histidine (D-his), allantoin. Growth was scored after 2 to 3 d of incubation at 30 °C.

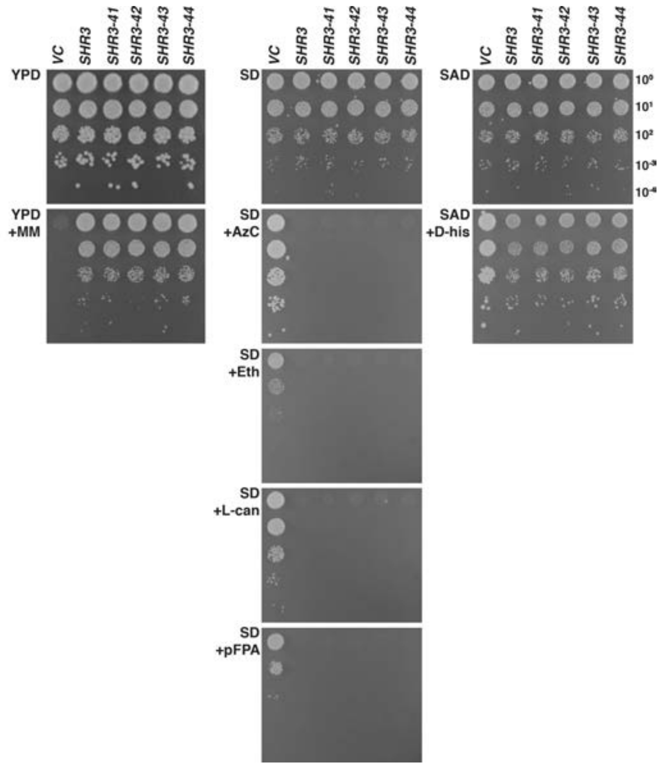


**Figure S3. Growth-based assessment of Shr3 substrate specificity - III**

Growth of serially diluted cell suspensions from strain JKY2 (*shr3*Δ) expressing indicated Shr3-mutant proteins spotted on indicated media. Concentrations of toxic amino analogues and nitrogen sources were as follows: 200 µg/ml metsulfuron-methyl (MM), yeast extract and peptone; 1 mM azetidine-2-carboxylate (AzC), ammonium; 10 µg/ml DL-ethionine (Eth), ammonium; 1 µg/ml L-canavanine (L-can), ammonium; 50 µg/ml p-Fluoro-DL-phenylalanine (pFPA), ammonium; 0.5% w/v D-histidine (D-his), allantoin. Growth was scored after 2 to 3 d of incubation at 30 °C.

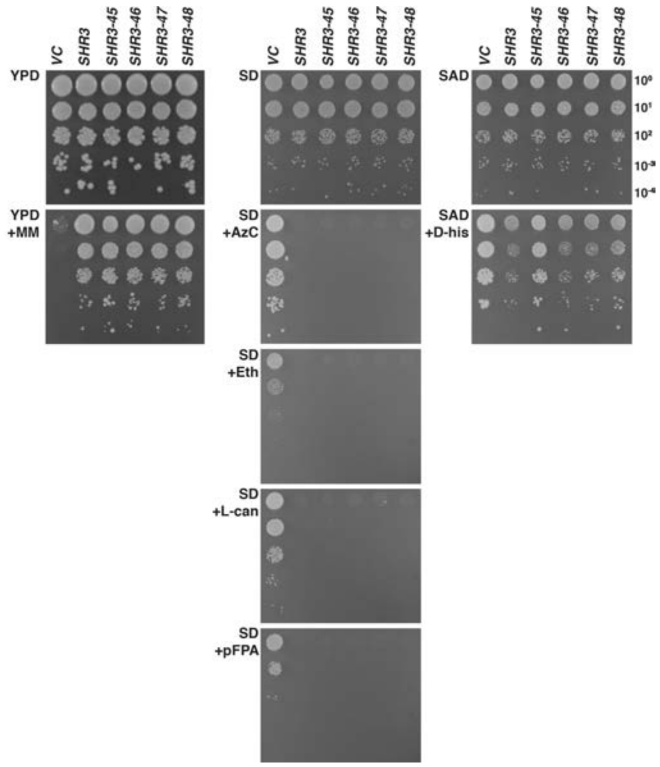


**Figure S4. Growth-based assessment of Shr3 substrate specificity - IV**

Growth of serially diluted cell suspensions from strain JKY2 (*shr3*Δ) expressing indicated Shr3-mutant proteins spotted on indicated media. Concentrations of toxic amino analogues and nitrogen sources were as follows: 200 µg/ml metsulfuron-methyl (MM), yeast extract and peptone; 1 mM azetidine-2-carboxylate (AzC), ammonium; 10 µg/ml DL-ethionine (Eth), ammonium; 1 µg/ml L-canavanine (L-can), ammonium; 50 µg/ml p-Fluoro-DL-phenylalanine (pFPA), ammonium; 0.5% w/v D-histidine (D-his), allantoin. Growth was scored after 2 to 3 d of incubation at 30 °C.

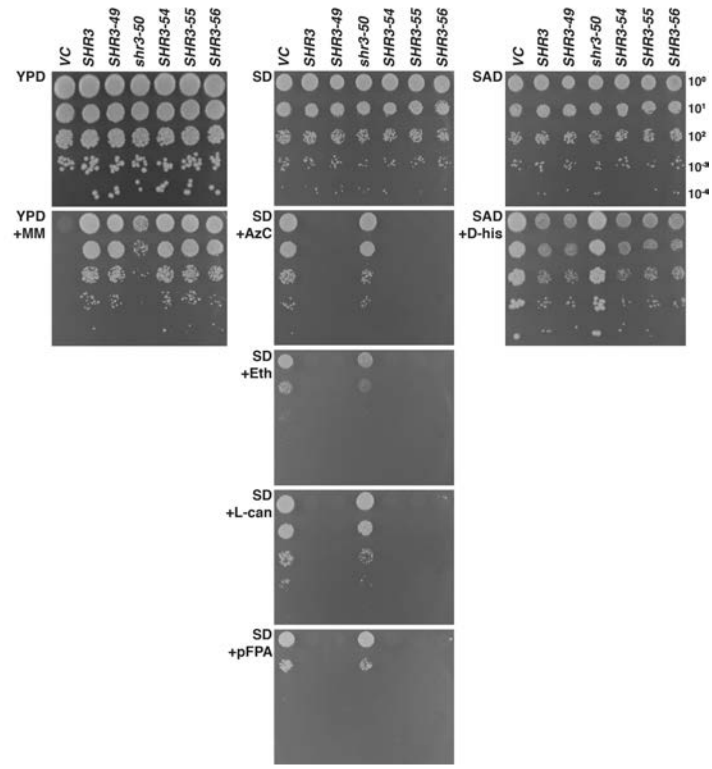


**Figure S5. Growth-based assessment of Shr3 substrate specificity - V**

Growth of serially diluted cell suspensions from strain JKY2 (*shr3*Δ) expressing indicated Shr3-mutant proteins spotted on indicated media. Concentrations of toxic amino analogues and nitrogen sources were as follows: 200 µg/ml metsulfuron-methyl (MM), yeast extract and peptone; 1 mM azetidine-2-carboxylate (AzC), ammonium; 10 µg/ml DL-ethionine (Eth), ammonium; 1 µg/ml L-canavanine (L-can), ammonium; 50 µg/ml p-Fluoro-DL-phenylalanine (pFPA), ammonium; 0.5% w/v D-histidine (D-his), allantoin. Growth was scored after 2 to 3 d of incubation at 30 °C.

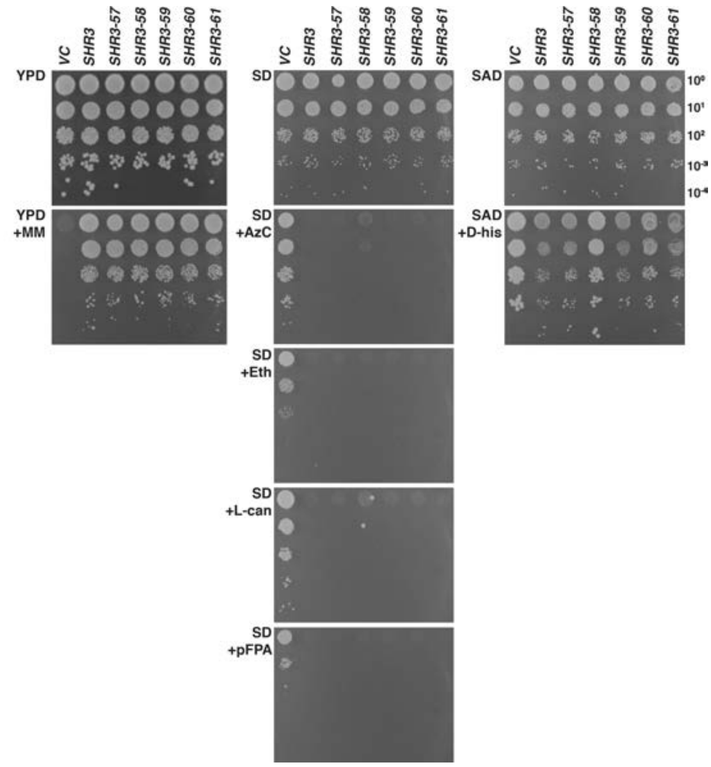


**Figure S6. Growth-based assessment of Shr3 substrate specificity - VI**

Growth of serially diluted cell suspensions from strain JKY2 (*shr3*Δ) expressing indicated Shr3-mutant proteins spotted on indicated media. Concentrations of toxic amino analogues and nitrogen sources were as follows: 200 µg/ml metsulfuron-methyl (MM), yeast extract and peptone; 1 mM azetidine-2-carboxylate (AzC), ammonium; 10 µg/ml DL-ethionine (Eth), ammonium; 1 µg/ml L-canavanine (L-can), ammonium; 50 µg/ml p-Fluoro-DL-phenylalanine (pFPA), ammonium; 0.5% w/v D-histidine (D-his), allantoin. Growth was scored after 2 to 3 d of incubation at 30 °C.

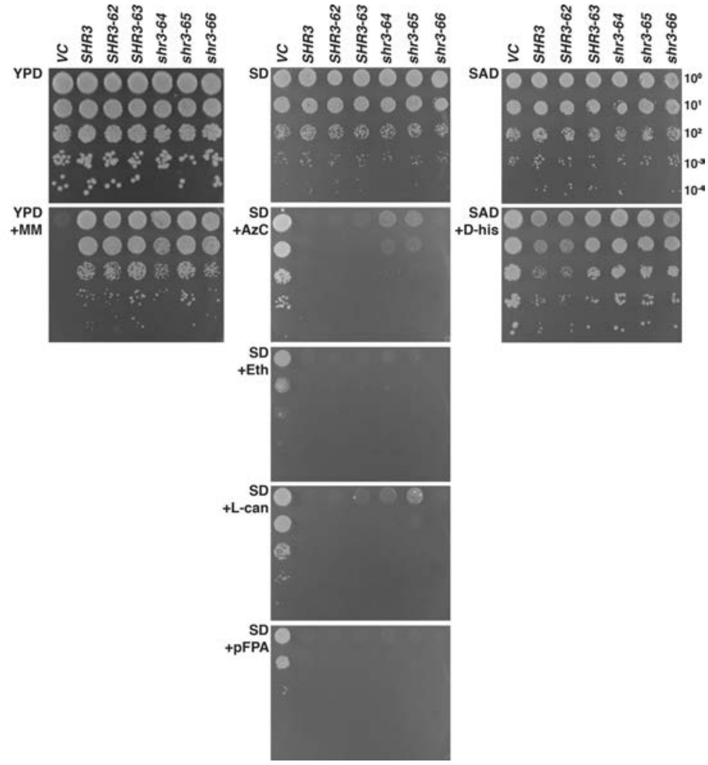


**Figure S7. Growth-based assessment of Shr3 substrate specificity - VII**

Growth of serially diluted cell suspensions from strain JKY2 (*shr3*Δ) expressing indicated Shr3-mutant proteins spotted on indicated media. Concentrations of toxic amino analogues and nitrogen sources were as follows: 200 µg/ml metsulfuron-methyl (MM), yeast extract and peptone; 1 mM azetidine-2-carboxylate (AzC), ammonium; 10 µg/ml DL-ethionine (Eth), ammonium; 1 µg/ml L-canavanine (L-can), ammonium; 50 µg/ml p-Fluoro-DL-phenylalanine (pFPA), ammonium; 0.5% w/v D-histidine (D-his), allantoin. Growth was scored after 2 to 3 d of incubation at 30 °C.

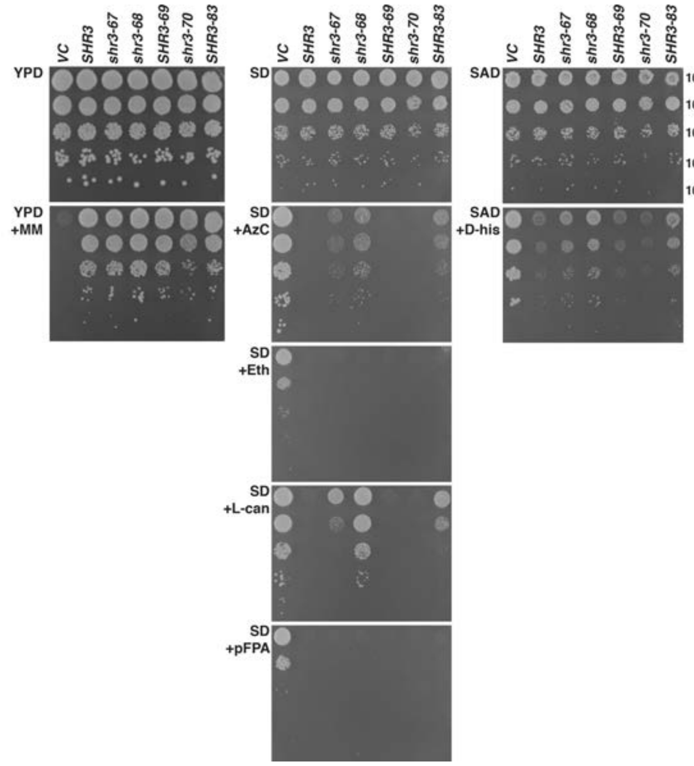


**Figure S8. Growth-based assessment of Shr3 substrate specificity - VIII**

Growth of serially diluted cell suspensions from strain JKY2 (*shr3*Δ) expressing indicated Shr3-mutant proteins spotted on indicated media. Concentrations of toxic amino analogues and nitrogen sources were as follows: 200 µg/ml metsulfuron-methyl (MM), yeast extract and peptone; 1 mM azetidine-2-carboxylate (AzC), ammonium; 10 µg/ml DL-ethionine (Eth), ammonium; 1 µg/ml L-canavanine (L-can), ammonium; 50 µg/ml p-Fluoro-DL-phenylalanine (pFPA), ammonium; 0.5% w/v D-histidine (D-his), allantoin. Growth was scored after 2 to 3 d of incubation at 30 °C.

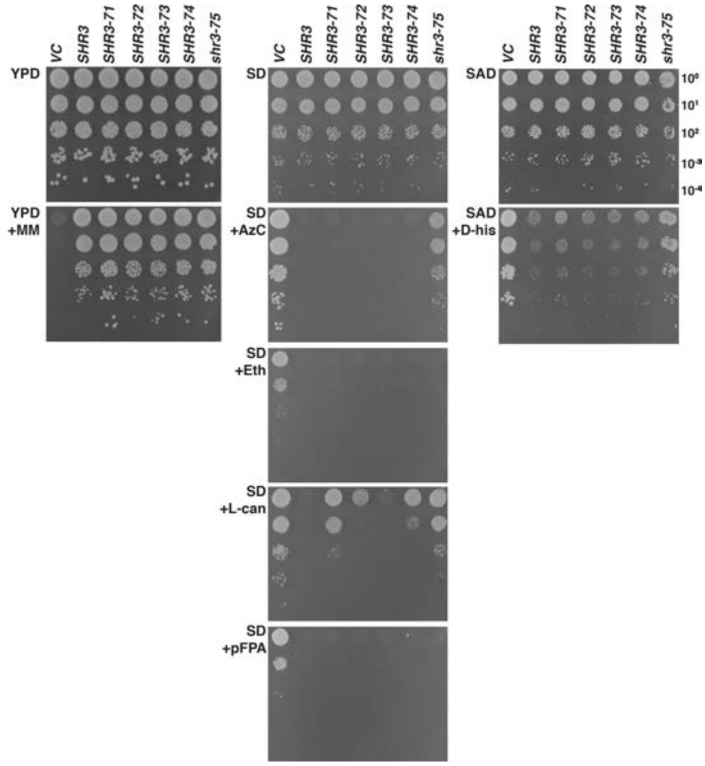


**Figure S9. Growth-based assessment of Shr3 substrate specificity - IX**

Growth of serially diluted cell suspensions from strain JKY2 (*shr3*Δ) expressing indicated Shr3-mutant proteins spotted on indicated media. Concentrations of toxic amino analogues and nitrogen sources were as follows: 200 µg/ml metsulfuron-methyl (MM), yeast extract and peptone; 1 mM azetidine-2-carboxylate (AzC), ammonium; 10 µg/ml DL-ethionine (Eth), ammonium; 1 µg/ml L-canavanine (L-can), ammonium; 50 µg/ml p-Fluoro-DL-phenylalanine (pFPA), ammonium; 0.5% w/v D-histidine (D-his), allantoin. Growth was scored after 2 to 3 d of incubation at 30 °C.

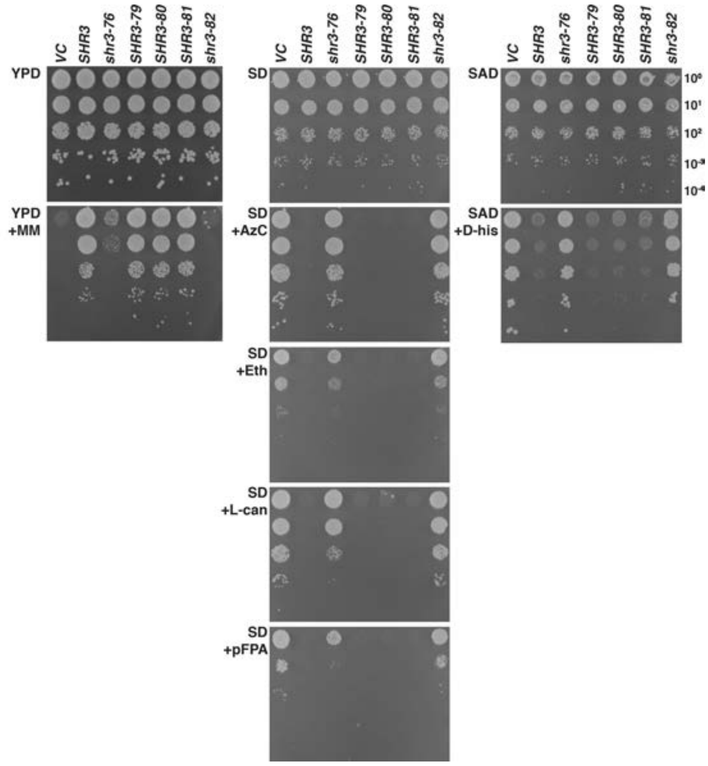


**Figure S10. Growth-based assessment of Shr3 substrate specificity - X**

Growth of serially diluted cell suspensions from strain JKY2 (*shr3*Δ) expressing indicated Shr3-mutant proteins spotted on indicated media. Concentrations of toxic amino analogues and nitrogen sources were as follows: 200 µg/ml metsulfuron-methyl (MM), yeast extract and peptone; 1 mM azetidine-2-carboxylate (AzC), ammonium; 10 µg/ml DL-ethionine (Eth), ammonium; 1 µg/ml L-canavanine (L-can), ammonium; 50 µg/ml p-Fluoro-DL-phenylalanine (pFPA), ammonium; 0.5% w/v D-histidine (D-his), allantoin. Growth was scored after 2 to 3 d of incubation at 30 °C.

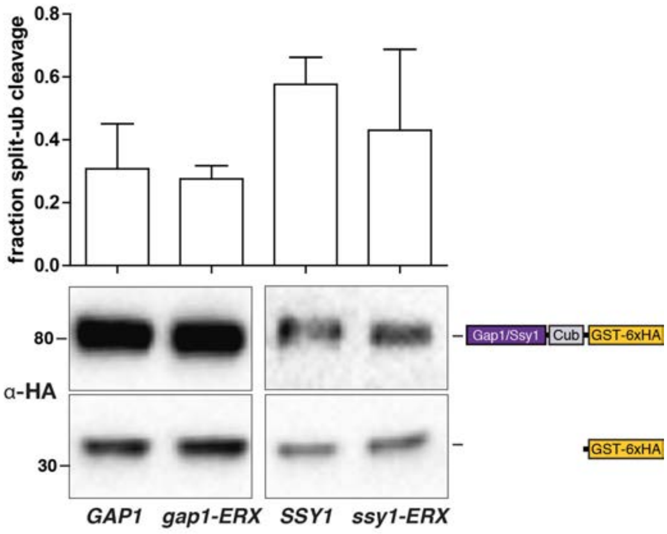


**Figure S11. Effect of ER exit motif mutations on Shr3-AAP interactions**

Strains FGY135 (*gap1Δ shr3Δ*) expressing *SHR3-NubA* (pPL1262) and carrying pPL1257 (*GAP1-Cub-GST*), pIM19 (*SSY1-Cub-GST*), pIM28 (*gap1-ERX$_{AAA}$-Cub-GST*) or pIM29 (*ssy1-ERX$_{AAA}$-Cub-GST*) were induced with 2% galactose for 1 h. Extracts were prepared, separated by SDS-PAGE and analyzed by immunoblotting using α-HA antibody. The signal intensities of the immunoreactive forms of full-length and cleaved Gap1 and Ssy1 constructs were quantified. The fraction of split-ubiquitin cleavage was determined; the mean values plotted with error bars showing standard deviation (n=3).

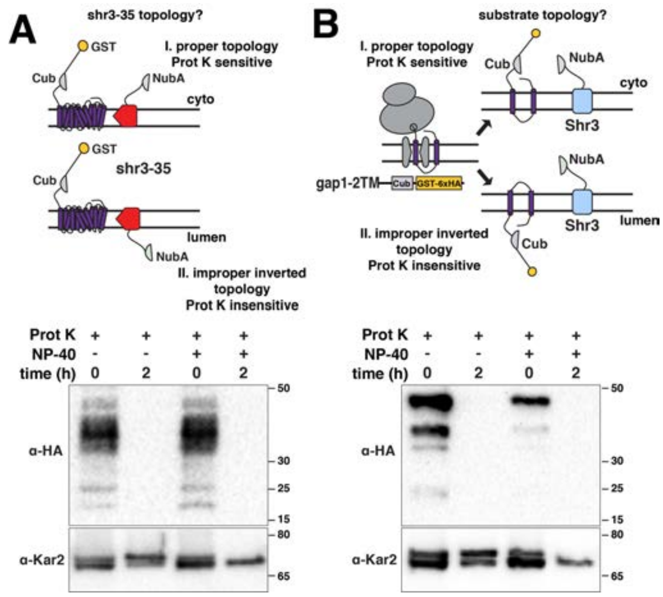


**Figure S12. Protease cleavage assay to assess the topology of shr3-35-NubA and gap1-2TM-Cub-GST**

(A) Schematic diagram of the split ubiquitin shr3-35-NubA construct (pAR67) and possible topological orientations: I. proper topology, i.e., TM in native orientation with NubA exposed to the cytoplasm and Proteinase K (Prot K) sensitive; II. improper inverted topology, i.e., TM in non-native orientation with NubA oriented towards the lumen and Prot K insensitive in the absence of detergent (NP-40). The ER-membrane topology of inserted shr3-35-NubA was examined in intact microsomes isolated from strain FGY135 (*gap1Δ shr3Δ*) carrying pPL1257 (*GAP1-Cub-GST*) and pAR67 (*shr3-35-NubA*). Microsomes were incubated with Prot K for 2 h with or without addition of 0.2% NP-40 as indicated. shr3-35 was detected with the use of a-Shr3 directed to a C-terminally located epitope. Kar2 served as a control for microsome integrity. (B) Schematic diagram of the split ubiquitin gap1-2TM-Cub-GST construct (pIM1) and possible topological orientations: I. proper topology, i.e., TM in native orientation with Cub-GST exposed to the cytoplasm and Proteinase K (Prot K) sensitive; II. improper inverted topology, i.e., TM in non-native orientation with Cub-GST oriented towards the lumen and Prot K insensitive in the absence of detergent (NP-40). The ER-membrane topology of inserted gap1-2TM-Cub-GST-6xHA was examined in intact microsomes isolated from strain HKY15 (*ssy1Δ*) carrying (*gap1-2TM-Cub-GST*) and incubated with Prot K for 2 h with or without addition of 0.2% NP-40 as indicated. Kar2 served as a control for microsome integrity.

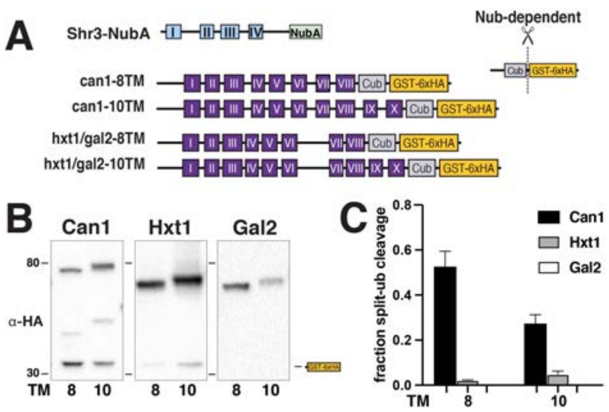


**Figure S13. Interactions between Shr3-NubA and can1-8TM, -10TM-Cub-GST**

(A) Schematic diagram of the split ubiquitin truncated constructs used to evaluate Shr3-can1 and Shr3-hxt1/gal2 interactions. (B) Strain FGY135 (*gap1*Δ *shr3*Δ) carrying pPL1262 (*SHR3-NubA*) and pIM46 (*can1-8TM-Cub-GST*), pIM47 (*can1-10TM-Cub-GST*), pIM37 (*hxt1-8TM-Cub-GST*), pIM38 (*hxt1-10TM-Cub-GST*), pIM43 (*gal2-8TM-Cub-GST*) or pIM44 (*gal2-10TM-Cub-GST*) was induced with 2% galactose for 1 h. Extracts were prepared, separated by SGS-PAGE and analyzed by immunoblotting using antibody against the 6xHA. (C) The signal intensities of the immunoreactive forms of uncleaved Cub constructs and cleaved interaction marker (GST-6xHA) were quantitated; the mean values of the fraction of split ubiquitin is plotted with error bars showing standard deviation (n=3). The quantitation corresponding to hxt1-8TM-Cub-GST, hxt1-10TM-Cub-GST, gal2-8TM-Cub-GST and gal2-10TM-Cub-GST are the same as presented in Fig. 6C. The immunoblot images corresponding to hxt1 and gal2 Cub-constructs are from independent biological replicates than depicted in Fig. 6B.

Table S1 Strains

| Strain | Genotype | Reference |
|---|---|---|
| JKY2 | *MATa ura3-52 shr3Δ6* | (Martins et al., 2019) |
| HKY15 | *MATa ura3-52 lys2Δ201 ssy1Δ12:: hisG-URA3-kanr-hisG* | (Forsberg and Ljungdahl, 2001) |
| FGY15 | *MATa ura3-52 lys2Δ201 leu2-3,112 gap1Δ2::LEU2* | (Gilstring and Ljungdahl, 2000) |
| FGY135 | *MATa ura3-52 lys2Δ201 leu2-3,112 gap1Δ2::LEU2 shr3Δ6* | (Gilstring and Ljungdahl, 2000) |

Table S2. Plasmids

| Plasmid | Description | Reference |
|---|---|---|
| pPL210 | pRS316 (URA3) containing *SHR3* | (Ljungdahl et al., 1992) |
| pJK92 | pRS317 (LYS2) containing *GAP1* | (Kota et al., 2007) |
| pCA204 | PRS316 (URA3) containing *STP1-13xMYC-kanMX* | (Andréasson et al., 2006) |
| pAR009 | pRS316 (URA3) containing *SHR3-30* mut aa 2-4 | This work |
| pAR010 | pRS316 (URA3) containing *SHR3-31* mut aa 5-7 | This work |
| pAR001 | pRS316 (URA3) containing *SHR3-32* mut aa 8-10 | This work |
| pAR002 | pRS316 (URA3) containing *SHR3-33* mut aa 11-13 | This work |
| pAR003 | pRS316 (URA3) containing *SHR3-34* mut aa 14-16 | This work |
| pAR004 | pRS316 (URA3) containing *shr3-35* mut aa 17-19 | This work |
| pPL1330 | pRS316 (URA3) containing *SHR3-36* mut aa 17-18 | This work |
| pAR047 | pRS316 (URA3) containing *shr3-37* mut aa 18-19 | This work |
| pAR037 | pRS316 (URA3) containing *SHR3-38* mut aa 17+19 | This work |
| pAR005 | pRS316 (URA3) containing *SHR3-39* mut aa 20-22 | This work |
| pAR006 | pRS316 (URA3) containing *SHR3-40* mut aa 23-25 | This work |
| pAR007 | pRS316 (URA3) containing *SHR3-41* mut aa 26-28 | This work |
| pAR008 | pRS316 (URA3) containing *SHR3-42* mut aa 29-31 | This work |
| pAR011 | pRS316 (URA3) containing *SHR3-43* mut aa 30-32 | This work |
| pAR012 | pRS316 (URA3) containing *SHR3-44* mut aa 33-35 | This work |
| pAR013 | pRS316 (URA3) containing *SHR3-45* mut aa 36-38 | This work |
| pAR014 | pRS316 (URA3) containing *SHR3-46* mut aa 39-41 | This work |
| pAR015 | pRS316 (URA3) containing *SHR3-47* mut aa 42-44 | This work |
| pAR016 | pRS316 (URA3) containing *SHR3-48* mut aa 45-47 | This work |
| pAR017 | pRS316 (URA3) containing *SHR3-49* mut aa 48-50 | This work |
| pAR018 | pRS316 (URA3) containing *shr3-50* mut aa 51-53 | This work |
| pAR051 | pRS316 (URA3) containing *SHR3-51* mut aa 51-52 | This work |
| pAR052 | pRS316 (URA3) containing *SHR3-52* mut aa 52-53 | This work |
| pAR050 | pRS316 (URA3) containing *shr3-53* mut aa 51+53 | This work |
| pAR019 | pRS316 (URA3) containing *SHR3-54* mut aa 54-56 | This work |
| pAR020 | pRS316 (URA3) containing *SHR3-55* mut aa 57-59 | This work |
| pAR021 | pRS316 (URA3) containing *SHR3-56* mut aa 60-62 | This work |
| pAR022 | pRS316 (URA3) containing *SHR3-57* mut aa 62-71 | This work |
| pPL1366 | pRS316 (URA3) containing *SHR3-58* mut aa 69-81 | This work |
| pAR023 | pRS316 (URA3) containing *SHR3-59* mut aa 82-84 | This work |
| pAR024 | pRS316 (URA3) containing *SHR3-60* mut aa 85-87 | This work |

| | | |
|---|---|---|
| pAR025 | pRS316 (URA3) containing *SHR3-61* mut aa 88-90 | This work |
| pAR026 | pRS316 (URA3) containing *SHR3-62*mut aa 91-93 | This work |
| pAR027 | pRS316 (URA3) containing *SHR3-63* mut aa 93-95 | This work |
| pPL1342 | pRS316 (URA3) containing *shr3-64* mut aa 96-101 | This work |
| pPL1343 | pRS316 (URA3) containing *SHR3-65* mut aa 100-107 | This work |
| pAR038 | pRS316 (URA3) containing *SHR3-66* mut aa 108-110 | This work |
| pAR039 | pRS316 (URA3) containing *SHR3-67*mut aa 111-113 | This work |
| PPL1346 | pRS316 (URA3) containing *SHR3-68* mut aa 114-119 | This work |
| pAR028 | pRS316 (URA3) containing *SHR3-69* mut aa 117-119 | This work |
| pAR029 | pRS316 (URA3) containing *SHR3-70* mut aa 120-122 | This work |
| pAR030 | pRS316 (URA3) containing *SHR3-71* mut aa 123-125 | This work |
| pAR031 | pRS316 (URA3) containing *SHR3-72* mut aa 126-128 | This work |
| pAR032 | pRS316 (URA3) containing *SHR3-73* mut aa 129-131 | This work |
| pAR033 | pRS316 (URA3) containing *SHR3-74* mut aa 132-133 | This work |
| pPL1348 | pRS316 (URA3) containing *SHR3-75* mut aa 134-138 | This work |
| pPL1349 | pRS316 (URA3) containing *shr3-76* mut aa 139-142 | This work |
| pAR048 | pRS316 (URA3) containing *shr3-77* mut aa 139-140 | This work |
| pAR049 | pRS316 (URA3) containing *SHR3-78* mut aa 141-142 | This work |
| pPL1351 | pRS316 (URA3) containing *SHR3-79* mut aa 145-153 | This work |
| pAR034 | pRS316 (URA3) containing *SHR3-80* mut aa 154-156 | This work |
| pPL1353 | pRS316 (URA3) containing *SHR3-81* mut aa 158-163 | This work |
| pPL1338 | pRS316 (URA3) containing *shr3-82* mut aa 31-41 | This work |
| pPL1347 | pRS316 (URA3) containing *SHR3-83* mut aa 120-123 | This work |
| pAR041 | pRS316 (URA3) containing *shr3Δ90* mut aa Δ34-48 | This work |
| pAR042 | pRS316 (URA3) containing *shr3Δ91* mut aa Δ39-47 | This work |
| pAR043 | pRS316 (URA3) containing *shr3Δ92* mut aa Δ44-54 | This work |
| pAR044 | pRS316 (URA3) containing *shr3Δ93* mut aa Δ55-60 | This work |
| pAR045 | pRS316 (URA3) containing *shr3Δ94* mut aa Δ121-127 | This work |
| pAR67 | pRS316 (URA3) containing *shr3-35-NubA* | This work |
| pAR76 | pRS316 (URA3) containing *SHR3Δ94-NubA* | This work |
| pPL1257 | pRS317 (LYS2) containing p*GAL-GAP1-Cub-GST-6xHA* | This work |
| pPL1262 | pRS316 (URA3) containing *SHR3-NubA* | This work |
| pIM1 | pRS317 (LYS2) containing p*GAL-gap1-TM2-Cub-GST-6xHA* | This work |
| pIM2 | pRS317 (LYS2) containing p*GAL-gap1-TM4-Cub-GST-6xHA* | This work |
| pIM3 | pRS317 (LYS2) containing p*GAL-gap1-TM6-Cub-GST-6xHA* | This work |
| pIM4 | pRS317 (LYS2) containing p*GAL-gap1-TM8-Cub-GST-6xHA* | This work |
| pIM5 | pRS317 (LYS2) containing p*GAL-gap1-TM10-Cub-GST-6xHA* | This work |
| pIM6 | pRS317 (LYS2) containing p*GAL-AGP1-Cub-GST-6xHA* | This work |
| pIM7 | pRS317 (LYS2) containing p*GAL-BAP2-Cub-GST-6xHA* | This work |
| pIM8 | pRS317 (LYS2) containing p*GAL-CAN1-Cub-GST-6xHA* | This work |
| pIM9 | pRS317 (LYS2) containing p*GAL-agp1-TM2-Cub-GST-6xHA* | This work |
| pIM10 | pRS317 (LYS2) containing p*GAL-agp1-TM4-Cub-GST-6xHA* | This work |
| pIM11 | pRS317 (LYS2) containing p*GAL-agp1-TM6-Cub-GST-6xHA* | This work |
| pIM12 | pRS317 (LYS2) containing p*GAL-agp1-TM8-Cub-GST-6xHA* | This work |
| pIM13 | pRS317 (LYS2) containing p*GAL-agp1-TM10-Cub-GST-6xHA* | This work |

| | | |
|---|---|---|
| pIM16 | pRS317 (LYS2) containing p*GAL-gap1-TM12-Cub-GST-6xHA* | This work |
| pIM17 | pRS317 (LYS2) containing p*GAL-GNP1-Cub-GST-6xHA* | This work |
| pIM18 | pRS317 (LYS2) containing p*GAL-LYP1-Cub-GST-6xHA* | This work |
| pIM19 | pRS317 (LYS2) containing p*GAL-SSY1-Cub-GST-6xHA* | This work |
| pIM20 | pRS317 (LYS2) containing p*GAL-ssy1-TM2-Cub-GST-6xHA* | This work |
| pIM21 | pRS317 (LYS2) containing p*GAL-ssy1-TM4-Cub-GST-6xHA* | This work |
| pIM22 | pRS317 (LYS2) containing p*GAL-ssy1-TM6-Cub-GST-6xHA* | This work |
| pIM23 | pRS317 (LYS2) containing p*GAL-ssy1-TM8-Cub-GST-6xHA* | This work |
| pIM24 | pRS317 (LYS2) containing p*GAL-ssy1-TM10-Cub-GST-6xHA* | This work |
| pIM25 | pRS317 (LYS2) containing p*GAL-ssy1-TM12-Cub-GST-6xHA* | This work |
| pIM26 | pRS317 (LYS2) containing p*GAL-agp1-TM12-Cub-GST-6xHA* | This work |
| pIM28 | pRS317 (LYS2) containing p*GAL-gap1-ERX-Cub-GST-6xHA* | This work |
| pIM29 | pRS317 (LYS2) containing p*GAL-ssy1-ERX-Cub-GST-6xHA* | This work |
| pIM32 | pRS317 (LYS2) containing p*GAL-HXT1-Cub-GST-6xHA* | This work |
| pIM33 | pRS317 (LYS2) containing p*GAL-GAL2-Cub-GST-6xHA* | This work |
| pIM34 | pRS317 (LYS2) containing p*GAL-hxt1-TM2-Cub-GST-6xHA* | This work |
| pIM35 | pRS317 (LYS2) containing p*GAL-hxt1-TM4-Cub-GST-6xHA* | This work |
| pIM36 | pRS317 (LYS2) containing p*GAL-hxt1-TM6-Cub-GST-6xHA* | This work |
| pIM37 | pRS317 (LYS2) containing p*GAL-hxt1-TM8-Cub-GST-6xHA* | This work |
| pIM38 | pRS317 (LYS2) containing p*GAL-hxt1-TM10-Cub-GST-6xHA* | This work |
| pIM39 | pRS317 (LYS2) containing p*GAL-hxt1-TM12-Cub-GST-6xHA* | This work |
| pIM40 | pRS317 (LYS2) containing p*GAL-gal2-TM2-Cub-GST-6xHA* | This work |
| pIM41 | pRS317 (LYS2) containing p*GAL-gal2-TM4-Cub-GST-6xHA* | This work |
| pIM42 | pRS317 (LYS2) containing p*GAL-gal2-TM6-Cub-GST-6xHA* | This work |
| pIM43 | pRS317 (LYS2) containing p*GAL-gal2-TM8-Cub-GST-6xHA* | This work |
| pIM44 | pRS317 (LYS2) containing p*GAL-gal2-TM10-Cub-GST-6xHA* | This work |
| pIM45 | pRS317 (LYS2) containing p*GAL-gal2-TM12-Cub-GST-6xHA* | This work |
| pIM46 | pRS317 (LYS2) containing *pGAL-can1-8TM-Cub-GST-6xHA* | This work |
| pIM47 | pRS317 (LYS2) containing *pGAL-can1-10TM-Cub-GST-6xHA* | This work |

**References**

Andréasson, C., Heessen, S., and Ljungdahl, P.O. (2006). Regulation of transcription factor latency by receptor-activated proteolysis. Genes Dev *20*, 1563-1568. 10.1101/gad.374206.

Forsberg, H., and Ljungdahl, P.O. (2001). Genetic and biochemical analysis of the yeast plasma membrane Ssy1p-Ptr3p-Ssy5p sensor of extracellular amino acids. Mol Cell Biol *21*, 814-826. 10.1128/MCB.21.3.814-826.2001.

Gilstring, C.F., and Ljungdahl, P.O. (2000). A method for determining the in vivo topology of yeast polytopic membrane proteins demonstrates that Gap1p fully integrates into the membrane independently of Shr3p. J. Biol. Chem. *275*, 31488-31495.

Kota, J., Gilstring, C.F., and Ljungdahl, P.O. (2007). Membrane chaperone Shr3 assists in folding amino acid permeases preventing precocious ERAD. The Journal of Cell Biology *176*, 617-628. 10.1083/jcb.200612100.

Ljungdahl, P.O., Gimeno, C.J., Styles, C.A., and Fink, G.R. (1992). SHR3: A novel component of the secretory pathway specifically required for localization of amino acid permeases in yeast. Cell *71*, 463-478. 10.1016/0092-8674(92)90515-E.

Martins, A., Ring, A., Omnus, D.J., Heessen, S., Pfirrmann, T., and Ljungdahl, P.O. (2019). Spatial and temporal regulation of the endoproteolytic activity of the SPS-sensor-controlled Ssy5 signaling protease. Mol Biol Cell *30*, 2709-2720. 10.1091/mbc.E19-02-0096.
